# Knowing what you don’t know: Estimating the uncertainty of feedforward and feedback inputs with prediction-error circuits

**DOI:** 10.1101/2023.12.13.571410

**Authors:** Loreen Hertäg, Katharina A. Wilmes, Claudia Clopath

## Abstract

At any moment, our brains receive a stream of sensory stimuli arising from the world we interact with. Simultaneously, neural circuits are shaped by feedback signals carrying predictions about the same inputs we experience. Those feedforward and feedback inputs often do not perfectly match. Thus, our brains have the challenging task of integrating these conflicting streams of information according to their reliabilities. However, how neural circuits keep track of both the stimulus and prediction uncertainty is not well understood. Here, we propose a network model whose core is a hierarchical prediction-error circuit. We show that our network can estimate the variance of the sensory stimuli and the uncertainty of the prediction using the activity of negative and positive prediction-error neurons. In line with previous hypotheses, we demonstrate that neural circuits rely strongly on feedback predictions if the perceived stimuli are noisy and the underlying generative process, that is, the environment is stable. Moreover, we show that predictions modulate neural activity at the onset of a new stimulus, even if this sensory information is reliable. In our network, the uncertainty estimation, and, hence, how much we rely on predictions, can be influenced by perturbing the intricate interplay of different inhibitory interneurons. We, therefore, investigate the contribution of those inhibitory interneurons to the weighting of feedforward and feedback inputs. Finally, we show that our network can be linked to biased perception and unravel how stimulus and prediction uncertainty contribute to the contraction bias.

## Introduction

To survive in an ever-changing environment, animals must flexibly adapt their behavior based on previously encoded and novel information. This adaptation is reflected in the information processing of neural networks underlying context-dependent behavior. For instance, when you walk down an unknown staircase in a fully lit basement, your brain might entirely rely on the feedforward (bottom-up) input your senses receive (Fig. 1A, left). In contrast, when you walk down the same stairs in complete darkness, your brain might rely entirely on feedback (top-down) signals generated from a staircase model it has formed over previous experiences (Fig. 1A, middle). But how do neural networks switch between a feedforward-dominated and a feedback-dominated processing mode? And how do neural networks in the brain combine both input streams wisely? For instance, if you hike down an unexplored mountain in very foggy conditions, your brain receives unreliable visual information. In addition, it can only draw on a shaky prediction about what to expect (Fig. 1A, right).

**Figure 1.**
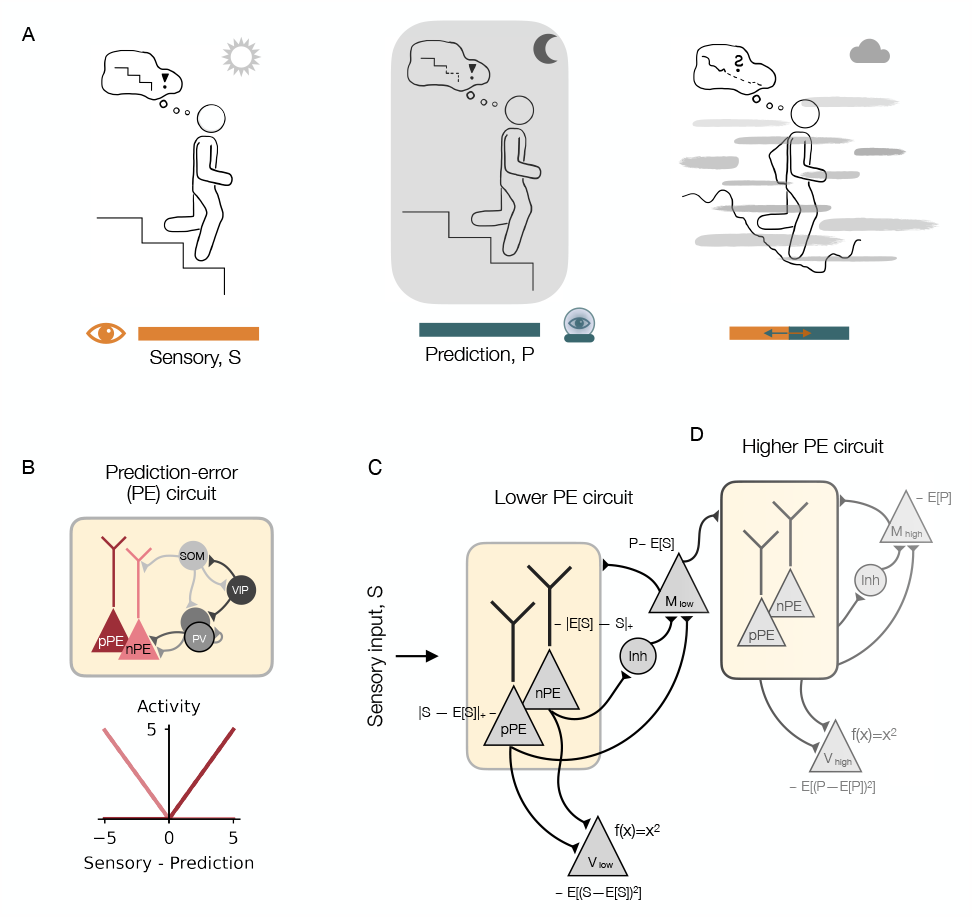
Neural network model to track both the uncertainty of sensory inputs and predictions. **(A)** Example illustration for context-dependent integration of information. Left: When walking down an unfamiliar staircase that is visible, the brain might rely solely on external sensory information. Middle: When walking down the same stairs without visual information, the brain might rely on predictions formed by previous experience. Right: When climbing down an unexplored mountain in foggy conditions, the brain might need to integrate sensory information and predictions simultaneously. **(B)** Top: Illustration of a prediction-error (PE) circuit with both negative and positive PE (nPE/pPE) neurons that receive inhibition from three different inhibitory interneuron types: parvalbumin-expressing (PV), somatostatin-expressing (SOM), and vasoactive intestinal peptide-expressing (VIP) interneurons. Local excitatory connections are not shown for clarity. Bottom: Responses of an nPE and pPE neuron. The nPE neuron only increases its activity relative to a baseline when the sensory input is weaker than predicted, while the pPE neuron only increases its activity relative to a baseline when the sensory input is stronger than predicted. **(C)** Illustration of network model that estimates the mean and variance of the external sensory stimuli. The core of this network model is the PE circuit shown in (B). The lower-level V neuron encodes the variance, while the lower-level M neuron encodes the mean of the sensory input. **(D)** Same as in (C) but the feedforward input is the activity of the lower-level M neuron.

A common hypothesis is that the brain weights different inputs according to their reliabilities. A prominent example of this hypothesis is Bayesian multisensory integration (see, e.g., Deneve and Pouget, 2004). According to this theory, neural networks represent information from multiple modalities by a linear combination of the uncertainty-weighted single-modality estimates. Multisensory integration is supported by several observations showing that animals can combine information from different modalities in a fashion that minimizes the variance of the final estimate (Ernst and Banks, 2002; Battaglia et al., 2003; Körding and Wolpert, 2004; Alais and Burr, 2004; Rowland et al., 2007; Gu et al., 2008; Fetsch et al., 2012). Here, we propose that the same concepts could be employed for the weighting of sensory inputs and predictions thereof (Körding and Wolpert, 2004; Yon and Frith, 2021). A central point in the weighting of inputs is the estimation of their variances as a measure of uncertainty. However, how the variance of both the sensory input and the prediction can be computed on the circuit level is not resolved yet.

We hypothesized that prediction error (PE) neurons provide the basis for the neural computation of variances. PEs are an integral part of the theory of predictive processing which states that the brain constantly compares incoming sensory information with predictions. If those predictions are wrong, the resulting PEs allow the network to revise the model of the world, thereby ensuring that the predictions become more accurate (Keller and Mrsic-Flogel, 2018). Experimental evidence suggests that these PEs may be represented in the activity of distinct groups of neurons, termed PE neurons (Eliades and Wang, 2008; Keller and Hahnloser, 2009; Ayaz et al., 2019; Audette et al., 2021). Moreover, these neurons may come in two types when excitatory neurons exhibit near-zero, spontaneous firing rates (Rao and Ballard, 1999; Keller and Mrsic-Flogel, 2018): negative PE (nPE) neurons only increase their activity when the prediction is *stronger* than the sensory input, while positive PE (pPE) neurons only increase their activity when the prediction is *weaker* than the sensory input. Indeed, it has been shown that excitatory neurons in rodent primary sensory areas can encode negative or positive PEs (Keller et al., 2012; Attinger et al., 2017; Jordan and Keller, 2020; Audette et al., 2021).

Here, we show that the unique response patterns of nPE and pPE neurons may provide the backbone for computing both the mean and the variance of sensory stimuli. Furthermore, we suggest a network model with a hierarchy of PE circuits to estimate the variance of the prediction, in addition to the variance of the sensory inputs. We show that in line with the ideas of multisensory integration, predictions are weighted more strongly than the sensory stimuli when the environment is stable (that is, predictable) and the sensory inputs are noisy. Moreover, we find that predictions are taken into account more at the beginning of a new trial than at the end, especially when the new sensory stimulus is reliable. In addition, we unravel the mechanisms underlying a neuromodulator-induced shift in the weighting of sensory inputs and predictions. In our model, these neuromodulators activate groups of inhibitory neurons such as parvalbumin-expressing (PV), somatostatin-expressing (SOM), and vasoactive intestinal peptide-expressing (VIP) interneurons (Markram et al., 2004; Rudy et al., 2011; Pfeffer et al., 2013; Jiang et al., 2015; Tremblay et al., 2016; Campagnola et al., 2022). These interneurons have been suggested to establish a multi-pathway balance of excitation and inhibition that is the basis for nPE and pPE neurons (Hertäg and Sprekeler, 2020; Hertäg and Clopath, 2022). By breaking this balance, the PE neurons change their baseline firing rate and gain, leading to a biased variance estimation. Finally, we show that this weighting can be understood as a neural manifestation of the contraction bias, that is, the magnitude of the represented sensory input is biased towards the mean of the past stimuli experienced (Hollingworth, 1910; Jazayeri and Shadlen, 2010; Ashourian and Loewenstein, 2011; Petzschner and Glasauer, 2011; Akrami et al., 2018; Meirhaeghe et al.).

## Results

### Prediction-error neurons as the basis for estimating the mean and variance of sensory stimuli

We hypothesize that the distinct response patterns of negative and positive prediction-error (nPE/pPE) neurons act as a backbone for estimating the mean and the variance of sensory stimuli. An nPE neuron only increases its activity relative to a baseline when the sensory input is weaker than predicted, while a pPE neuron only increases its activity relative to a baseline when the sensory input is stronger than predicted. Moreover, both nPE and pPE neurons remain at their baseline activities when the sensory input is fully predicted (Fig. 1B). If the prediction equals the mean of the sensory stimulus, the PE neurons, hence, encode the deviation from the mean. Thus, the squared sum of nPE and pPE neuron activity represents the variance of the feedforward input (provided that the PE neurons are silent without sensory stimulation).

To test our hypothesis, we study a rate-based mean-field network: the core network is a prediction-error (PE) circuit with excitatory nPE and pPE neurons, as well as inhibitory parvalbumin-expressing (PV), somatostatin-expressing (SOM), and vasoactive intestinal peptide-expressing (VIP) interneurons (Fig. 1B). While the excitatory neurons are simulated as two coupled point compartments to emulate the soma and dendrites of elongated pyramidal cells, respectively, all inhibitory cell types were modeled as point neurons. The connectivity of and inputs to the network were chosen such that the excitatory (E) and inhibitory (I) pathways onto the pyramidal cells were partially balanced. This balance that is only temporarily broken during mismatches has been shown to be necessary for nPE and pPE neurons to emerge (Hertäg and Sprekeler, 2020; Hertäg and Clopath, 2022, see Methods).

We assume that this core circuit is shaped by feedback connections (Larkum, 2013; Harris and Shepherd, 2015) that have been hypothesized to carry information about expectations or predictions (Mumford, 1992; Larkum, 2013; Friston, 2008). To account for predictions, we model a memory (M) neuron that integrates the activity of the PE neurons (Fig. 1C). Following Keller and Mrsic-Flogel (2018), we assume that the pPE neuron excites the memory neuron, while the nPE neuron inhibits this neuron (for instance, through lateral inhibition, here not modeled explicitly). Because feedback connections are shown to target the apical dendrites of pyramidal cells (Larkum, 2013) and interneurons located in superficial layers of the cortex (see, e.g. Tremblay et al., 2016), the memory neuron makes connections with the dendritic compartment of the PE neurons and some of the interneurons (here, VIP and PV neurons, see Methods for more details). We, furthermore, simulate a downstream neuron (termed V neuron), modeled as a leaky integrator with a quadratic activation function, that receives excitatory synapses from the PE neurons. In this setting, the V neuron encodes the variance of the sensory stimuli (see the lower-level subnetwork in Fig. 1C, the higher-level circuit is described later).

To show that this network can indeed represent the mean and the variance in the respective neurons, we stimulate it with a sequence of step-wise constant inputs drawn from a uniform distribution (Fig. 2A). We, hence, assume that the sensory stimulus varies over time. In line with the distinct response patterns for nPE and pPE neurons, these neurons change only slightly with increasing stimulus mean but increase strongly with input variance (Fig. 2B). In contrast, the three interneurons strongly increase with stimulus mean and only moderately increase with stimulus variance (Fig. 2C). The activity of the memory neuron gradually approaches the mean of the sensory inputs (Fig. 2D, middle), while the activity of the V neuron approaches the variance of those inputs (Fig. 2E, middle). We show that this holds for a wide range of input statistics (Fig. 2D-E, right) and input distributions (Fig. S1). Small deviations from the true mean occur mainly for large input variances, while the estimated variance is fairly independent of the input statistics tested.

**Figure 2.**
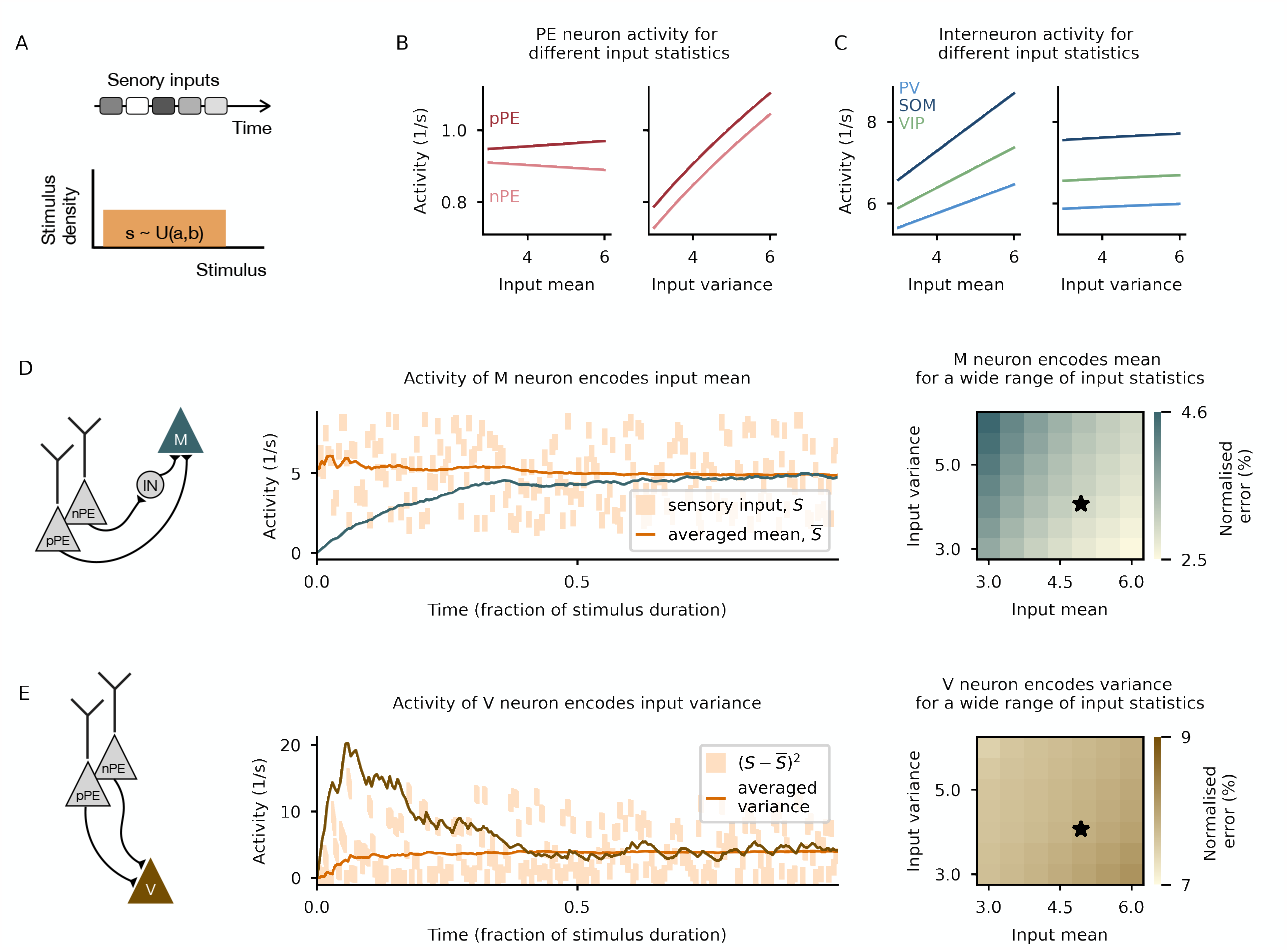
Prediction-error neurons as the basis for estimating mean and variance of sensory stimuli. **(A)** Illustration of the inputs with which the network (Fig. 1C) is stimulated. Network is exposed to a sequence of constant stimuli drawn from a uniform distribution. The gray shaded boxes symbolize different values from the distribution. **(B)** PE neuron activity hardly changes with stimulus strength (left) but strongly increases with stimulus variability (right). **(C)** Interneuron activity strongly changes with stimulus strength (left) but hardly changes with stimulus variability (right). **(D)** M neuron correctly encodes the mean of the sensory stimuli. Left: Illustration of the input synapses onto the M neuron. Middle: Activity of the M neuron over time for one example distribution (black star in right panel). Right: Normalised absolute difference between the averaged mean and the activity of the M neuron in the steady state for different parametrizations of the stimulus distribution. **(E)** V neuron correctly encodes the variance of the sensory stimuli. Left: Illustration of the input synapses onto the V neuron. Middle: Activity of the V neuron over time for one example distribution (black star in right panel). Right: Normalised absolute difference between the averaged variance and the activity of the V neuron in the steady state for different parametrizations of the stimulus distribution.

We verified our results in a heterogeneous network in which a population of neurons belongs to each neuron type of the PE circuit, and the synaptic connection strengths from each PE neuron onto the M and V neuron are different (see Methods, Fig. S2A). As before, the network can correctly estimate the mean and the variance of the sensory stimuli (Fig. S2B). Furthermore, we show that the errors with which the M and V neurons encode the stimulus statistics are independent of uncorrelated modulations of those connection strengths (Fig. S2C) and the sparsity of the network (Fig. S2E). When all connection strengths are collectively shifted to higher values, the error increases for the variance neuron, while it remains unaffected for the memory neuron.

While our mean-field network was designed to track the mean and the variance of stimuli that vary in time, we reasoned that the same principles apply to stimuli that vary across space. To show that, we simulated a population network that consists of unconnected replicates of the mean-field network described above (Fig. S3A). Each mean-field network receives a short, constant input from a different part of the receptive field. If the connection strengths from the PE neurons to the M and V neurons are adjusted accordingly (see Methods), the network correctly estimates the stimulus average and spatial uncertainty (Fig. S3B-C).

In summary, nPE and pPE neurons can serve as a basis to estimate the mean and the variance of sensory stimuli which vary over time and space.

### Estimating the uncertainty of both the sensory input and the prediction requires a hierarchy of PE circuits

Following the ideas of Bayesian multisensory integration, the weighting of sensory stimuli and predictions would require knowledge about their uncertainties. As we have shown in the previous section, the variance of the sensory stimulus can be estimated using PE neurons. We hypothesize that the same principles apply to computing the variance of the prediction. To show this, we augment the network with a *higher* PE circuit that receives feedforward synapses from the memory (M) neuron of the *lower* PE circuit (Fig. 1D). Both subnetworks are identical except for the M neuron in the higher PE circuit which is modeled with slower dynamics than the one in the lower PE circuit.

To test the network’s ability to estimate the variances correctly, we stimulated the network with a sequence of inputs whose mean can vary from trial to trial. More precisely, in each trial, the network is given a stimulus that is composed of *N*_in_ constant values, each drawn from a normal distribution and presented for *N*_step_ consecutive time steps. The variance of this distribution represents the stimulus noise. To account for potential changes in the environment, we draw the stimulus mean from a uniform distribution (Fig. 3A). Hence, the inputs change on two different time scales.

**Figure 3.**
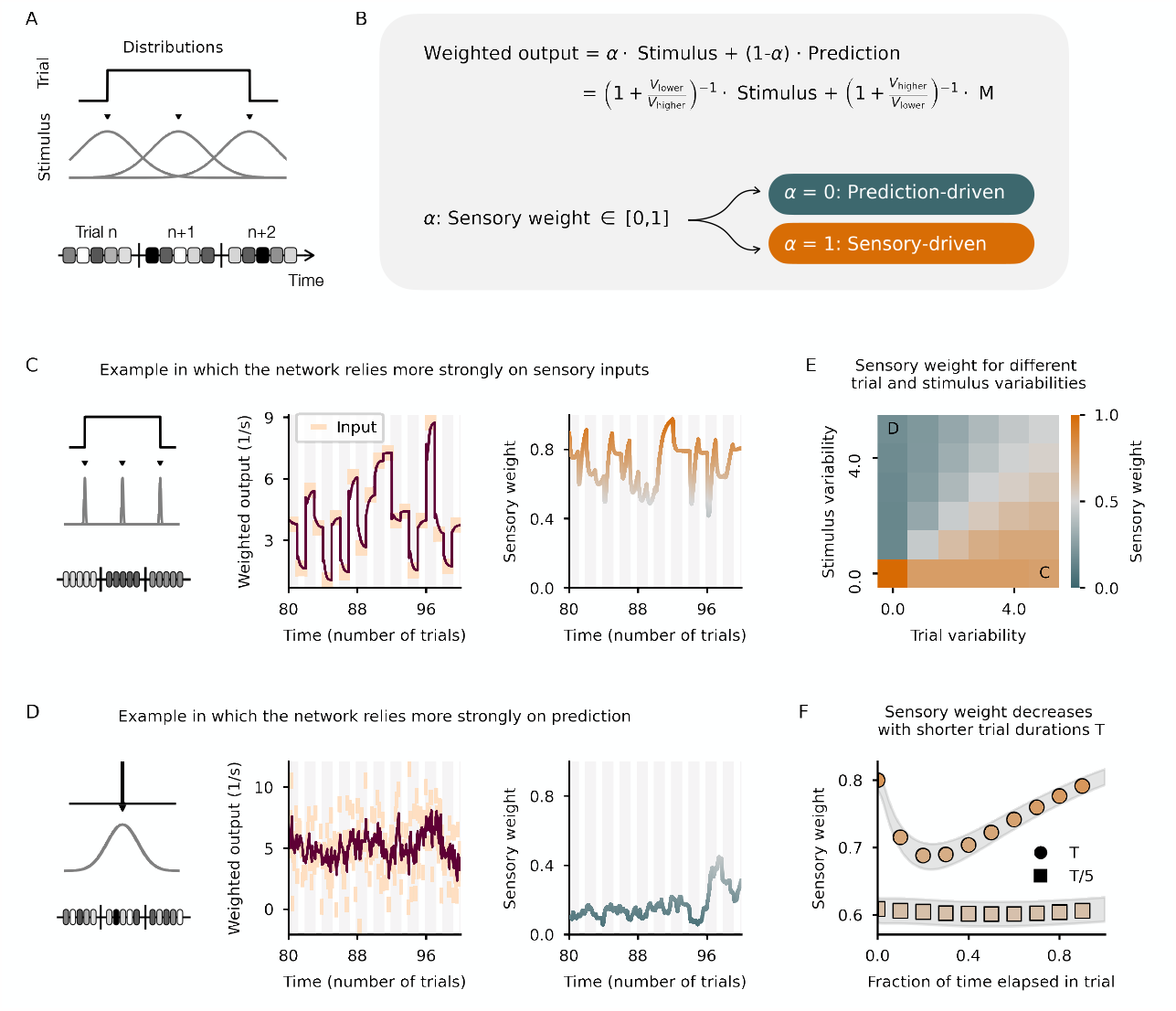
Estimating the uncertainty of both the sensory input and the prediction. **(A)** Illustration of the stimulation protocol. The network is exposed to a sequence of stimuli (one stimulus per trial). To account for stimulus variability, each stimulus is represented by 10 stimulus values drawn from a normal distribution. To account for the volatility of the environment, in each trial, the stimulus mean is drawn from a uniform distribution (denoted trial-to-trial variability). **(B)** Illustration of how the weighted output is calculated. The sensory weight *α* lies between zero (system relies perfectly on prediction) and one (system relies solely on the sensory input). **(C)** Limit case example in which the stimulus variability is zero but the trial-to-trial variability is high. Left: Illustration of the stimulation protocol. Middle: Weighted output follows closely the sensory stimuli. Right: Sensory weight (function of the variances, see B) close to 1, indicating that the network ignores the prediction. Input statistics shown in E. **(D)** Limit case example in which the stimulus variability is high but the trial-to-trial variability is zero. Left: Illustration of the stimulation protocol. Middle: Weighted output pushed towards the mean of the sensory stimuli. Right: Sensory weight close to zero, indicating that the network ignores the sensory stimuli. Input statistics shown in E. **(E)** Sensory weight for different input statistics. Predictions are weighted more strongly when the stimulus variability is larger than the trial-to-trial variability. **(F)** Sensory weight, averaged over many trials, for two different trial durations. Gray shading denotes the SEM. Predictions are weighted more strongly at the beginning of a new trial.

Following the formalism of multisensory integration (see, e.g. Pouget et al., 2013), we assume that the network’s output is a weighted sum of the feedforward sensory input and the feedback prediction. The weights assigned to each input stream are functions of the uncertainties, that is, the activities of the V neurons. The sensory weight captures how much the network relies on the sensory input (Fig. 2B). To test our network, we first consider two limit cases. In the first limit case, we show a low-variance stimulus that differs in each trial (low stimulus uncertainty, high trial-to-trial uncertainty, see Fig. 3C, left). According to the theory, the network should follow the sensory inputs closely and ignore the predictions. When we arithmetically calculate the weighted output (Fig. 3C, middle) and the sensory weight (Fig. 3C, right), the network indeed shows a clear preference for the sensory input. In the second limit case, we show a high-variance stimulus, the mean of which does not change from trial to trial (high stimulus uncertainty, low trial-to-trial uncertainty, see Fig. 3D, left). According to the theory, the network should downscale the sensory feedforward input and weight the prediction more strongly. Indeed, the weighted output of the network shows a clear tendency to the mean of the stimuli (Fig. 3D, middle), also reflected in the low sensory weight (Fig. 3D, right).

To validate the network responses fully, we systematically varied the trial and stimulus variability independently. If both variances are similar, the sensory weight approaches *0*.*5*, reflecting equal contribution of the sensory input and the prediction to the weighted output. Only if both variances are zero, the network represents the sensory input perfectly. In line with the limit case examples above, if the stimulus variance is larger than the trial variance, the network weights the prediction more strongly than the sensory input (Fig. 3E). Because the network dynamically estimates the sensory and prediction uncertainty, the sensory weight changes when the input statistics shifts (Fig. S4).

Inspecting closely the dynamics of our network, we noticed that the prediction is typically weighted higher at the beginning of a new trial than in the steady state. This is particularly pronounced in a sensory-driven input regime (see Fig. 3C). This is further confirmed in simulations in which the trail duration was shortened (Fig. 3F). Our model makes therefore the following experimentally testable prediction: sensory predictions influence neural activity more significantly in experiments that rely on fast stimulus changes.

It has been hypothesized, that some symptoms in psychiatric disorders like autism and schizophrenia can be ascribed to a pathological weighting of sensory inputs and predictions (Yon and Frith, 2021). We thus wondered which network properties might bias the estimation of the variances, and, consequently, the weighting of different input streams. We identified the time scales at which the memory neurons incorporate new information as a decisive factor in the integration of inputs. To show this, we varied the weights from the PE neurons onto the lower-level memory neuron. If the weights are too small (the memory neuron updates too slowly), the system relies too much on feedback predictions. In contrast, if the weights are too large (the memory neuron updates too fast), the system relies too much on the feedforward sensory information (Fig. S5A). While the speeds at which the activity of the memory neurons evolve influence the weighting of inputs, the precise activation function of the variance neurons is less pivotal. When we replaced the quadratic activation function with a linear, rectified function, the V neurons did not encode the variance but the average absolute deviation of the sensory stimuli. However, the sensory weight is only slightly shifted to larger values for low trial/high stimulus variability (Fig. S5B).

In summary, we show that the variances of both the sensory inputs and predictions thereof can be dynamically computed in networks comprising a lower and higher PE circuit. In such a network, predictions are given more weight at the beginning of a new stimulus, and if the sensory inputs are noisy while the environment is stable.

### Biasing the weighting of sensory inputs and predictions by neuromodulators

The brain’s flexibility and adaptability are supported by a plethora of neuromodulators which influence the activity of neurons in a variety of ways (Avery and Krichmar, 2017). A prominent target of neuromodulatory inputs is inhibitory neurons (Cardin, 2019; Hattori et al., 2017; Swanson and Maffei, 2019). Moreover, distinct interneuron types are differently (in-)activated by those neuromodulators (Wester and McBain, 2014; Hattori et al., 2017; Swanson and Maffei, 2019). We, therefore, wondered if and how the weighting of sensory inputs and predictions thereof may be biased when neuromodulators activate distinct interneuron types.

To this end, we modeled the presence of a neuromodulator by injecting an additional excitatory input into an interneuron type (while a neuromodulator can also suppress neuronal activity, we focus on the more common excitatory effects that have been described). We reasoned that the network effect of a neuromodulator not only depends on the interneuron type it targets but also on the inputs this neuron receives and the connections it makes with other neurons in the network. We, therefore, tested different mean-field networks that differ in the distribution of sensory inputs and predictions onto the interneurons, and the underlying connectivity. The commonality across those networks is that they exhibit an E/I balance of excitatory and inhibitory pathways onto the PE neurons (Hertäg and Clopath, 2022).

Across the different mean-field networks tested, increasing the activity of PV neurons biases the network’s output toward predictions (Fig. 4A left). In contrast, increasing VIP activity forces the networks to weigh both inputs more equally. As a consequence, predictions are overrated in a sensory-driven input regime, and, sensory inputs are overrated in a prediction-driven input regime (Fig. 4A right). Increasing SOM neuron activity, while qualitatively similar to increasing VIP neuron activity, depends on the mean-field network tested and the strength of activation (Fig. 4A middle).

**Figure 4.**
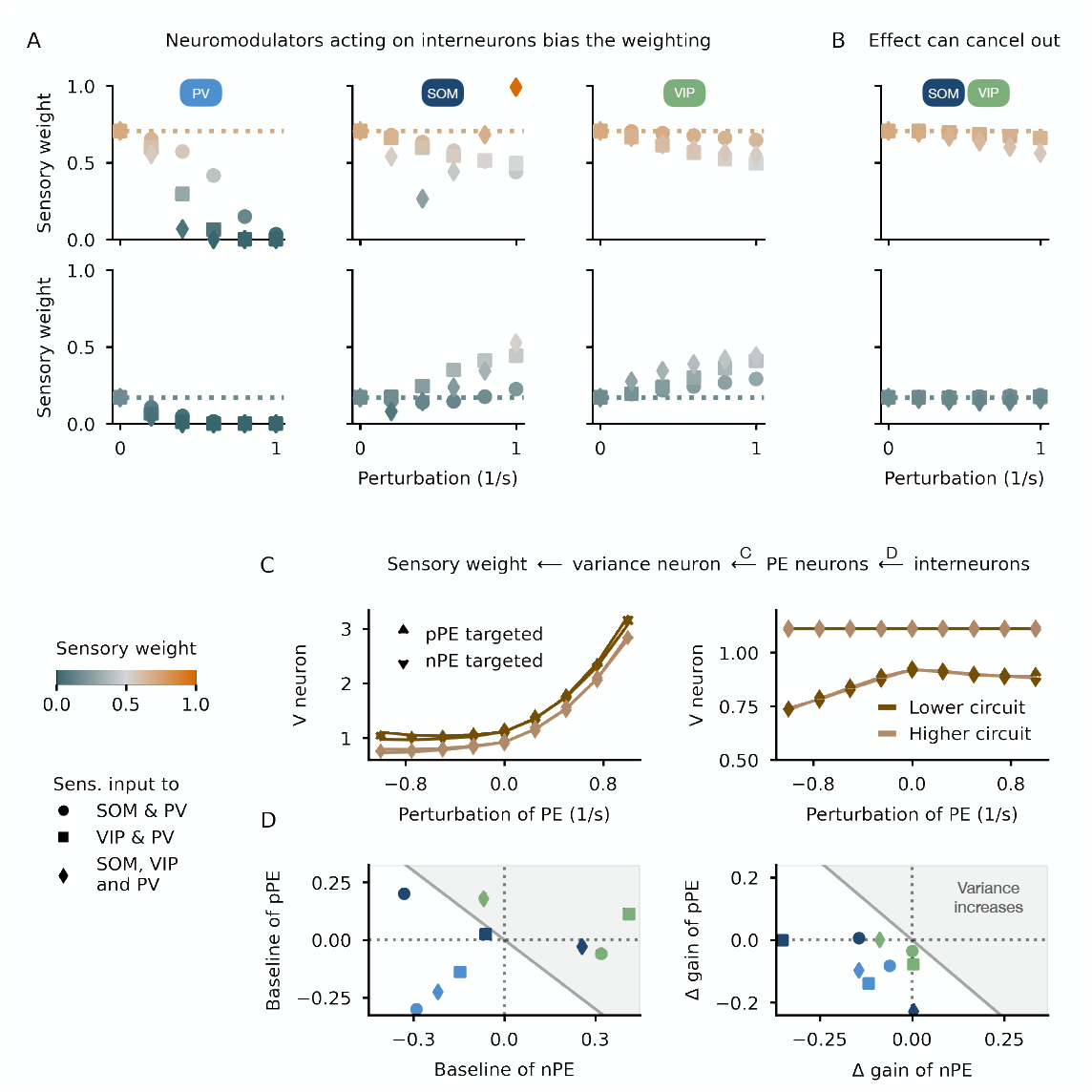
Neuromodulator-based shifts in the weighting of sensory inputs and predictions. **(A)** Neuromodulators acting on the interneurons can shift the weighting of sensory inputs and predictions. The changes depend on the type of interneuron targeted and the modulation strength (here simulated through an additional excitatory input). Considered are two limit cases (upper row: more sensory-driven before modulation, lower row: more prediction-driven before modulation). The results are shown for three different PE circuits (denotes by different markers). **(B)** When SOM and VIP neurons are equally modulated, the sensory weight remains unaffected. **(C)** The V neurons’ activities depend on the PE neurons. Hence, perturbing the nPE and pPE neuron changes the uncertainty estimation. Left: stimulating the pPE (triangle) or nPE (upside down triangle) affects the V neuron of the same subnetwork, denoted by matching marker and line colors (dark brown: lower circuit, light brown: higher circuit). Right: while stimulating the lower PE neurons affects the higher-order V neuron, stimulating the higher-order PE neurons does not change the activity of the lower-order V neuron (denoted by not matching marker and line colors). **(D)** The V neuron activity, and hence the sensory weight, changes as a result of the modulated PE neuron activity. The PE neuron activity, on the other hand, changes as a result of the interneurons being modulated. The interneurons change the baseline (left) and the gain (right) of the PE neurons. Whether an interneuron increases or decreases the estimated variance depends on both factors.

Neuromodulators are most likely increasing the activity of more than one interneuron type. To account for the co-activation of interneurons, we injected an excitatory input into two interneuron types at the same time and varied the strength with which each interneuron was modulated (Fig. S6). If PV neurons are the major target of a neuromodulator, the network is still biased toward predictions. If SOM and VIP neurons are equally stimulated, the weighting of sensory inputs and predictions remains largely unaffected (Fig. 4B), suggesting that the individual effects cancel out.

What are the network mechanisms underlying these observations? The sensory weight is a function of the lower and higher variance (V) neuron activity. Hence, any changes to the sensory weight result from changes to the neurons encoding the variances. In our network, the V neurons only receive excitatory synapses from PE neurons. As a consequence, any changes in the sensory weights upon activation of interneurons must be due to changes in the PE neurons. To disentangle the effect of nPE and pPE neurons, we perturbed those neurons individually in both the lower or higher subnetwork by injecting either an inhibitory or excitatory additional input (Fig. 4C). Stimulating either PE neuron in the lower subnetwork increases the activity of the lower-level V neuron strongly. Moreover, the higher-level V neuron is also slightly affected. This is because the lower-level memory neuron is also modulated by the lower-level PE neurons and makes feedforward connections to the higher-level PE circuit. In contrast, stimulating either PE neuron in the higher subnetwork increases the activity of the higher-level V neuron but leaves the lower-level neurons unaffected.

This suggests that to understand the effect of neuromodulators on the sensory weight, we need to unravel the effect of interneuron activation on PE neurons. Increasing interneuron activity leads to changes in the baseline and gain of PE neurons that bias the estimation of mean and variance (Fig. S8, see Methods). In all three networks tested, activating PV neurons decreases both baseline and gain of the PE neurons, leading to a decrease in the estimated variance (Fig. 4D & Fig. S9). Stimulating the SOM or VIP neuron decreases the gain in either nPE or pPE neuron. However, the baseline of those neurons can either decrease or increase depending on the connectivity with other neurons in the network. The summed effect over nPE and pPE neuron (Fig. S9) suggests that whether the activity of the V neuron increases or decreases depends on the input statistics: for low-mean stimuli, the elevated baseline activity dominates the changes in the variance, while for high-mean stimuli the changes in the gain dominate.

Altogether, we show that neuromodulators increasing the activity of interneurons bias the weighting of sensory inputs and predictions by changing the gain and baseline of PE neurons. Whether the sensory weight increases or decreases depends not only on the interneuron it targets but also on the network it is embedded in and the input regime.

### Explaining the contraction bias with the weighting of sensory inputs and predictions

We hypothesized that the weighted integration of sensory inputs and predictions thereof manifests in all-day behavior, in the form of a phenomenon called *contraction bias*. The contraction bias describes the tendency to overestimate sensory stimuli drawn from the lower end of a stimulus distribution and to underestimate stimuli drawn from the upper end of the same distribution. This *bias toward the mean* has been reported in different species and modalities (Hollingworth, 1910; Jazayeri and Shadlen, 2010; Ashourian and Loewenstein, 2011; Petzschner and Glasauer, 2011; Akrami et al., 2018; Meirhaeghe et al.).

We first investigated whether the network’s output can be interpreted as a neuronal manifestation of the contraction bias (see Methods for an illustrative analysis). To this end, we define the contraction bias as the trial-averaged difference between the weighted output and the sensory stimulus. When plotted over the trail-averaged stimuli, the bias is positive for stimuli below the mean of the input distribution and negative for stimuli above the mean (Fig. 5A), in line with a *bias toward the mean*. To measure the amount of bias in the network, we use the slope of the linear fit to the relationship between bias and trial stimulus. The larger the absolute slope, the larger the bias.

**Figure 5.**
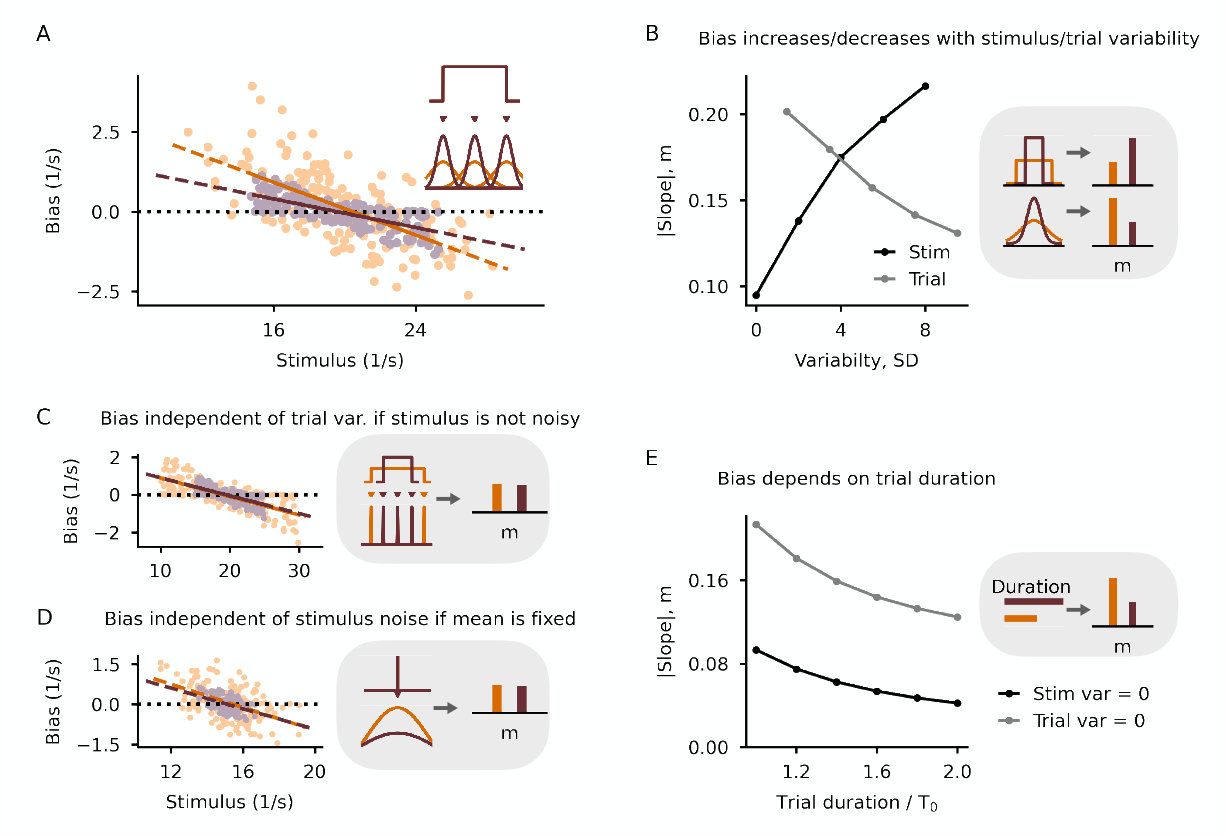
Mechanisms underlying the contraction bias. **(A)** Contraction bias in the model for two different stimulus uncertainties depicted in the inset. Bias is defined as the weighted output minus the stimulus mean. The absolute value of the slope of the linear fit, *m*, is a measure of the bias. The larger the slope, the larger the bias. **(B)** As a consequence of the sensory weight, the slope increases with stimulus variability (bias increases) and decreases with trial-to-trial variability (bias decreases). **(C)** Bias is independent of the trial-to-trial variability when the stimulus variability is zero. **(D)** Bias is independent of the stimulus variability when the trial-to-trial variability is zero. **(E)** The slope depends on the trial duration.

What are the underlying network factors that contribute to the neuronal contraction bias in the network? We have seen that how much the prediction is taken into account is determined by both the lower and higher-level V neurons encoding the variance of the stimulus and the prediction. Hence, the bias must be similarly influenced by these factors. When we increase the stimulus uncertainty, the bias increases (Fig. 5B). In contrast, when we increase the trial-to-trial uncertainty, the bias decreases (Fig. 5B).

To further disentangle the different sources of the bias, we first simulated a network without stimulus uncertainty (variance set to zero) for two trial-to-trail variances (volatility of the environment). In this case, the emerging contraction bias is independent of the volatility of the environment (Fig. 5C). We show mathematically that the bias results from the network output not yet reaching its new steady state within the trial duration (see Methods). In other words, the bias is the difference between the weighted output at the end of the trial and its steady state (the shown stimulus). How fast the new steady state is reached depends only on the time constants in the network and not the trial-to-trial variability. We next resume the limit case in which the stimulus uncertainty is high while the trial-to-trial uncertainty is zero. In this case, the contraction bias is also largely independent of the stimulus variance (Fig. 5D). Our mathematical analysis reveals that the bias is well described by the difference between the prediction, that is, mean stimulus over the history of all stimuli shown, and the current stimulus, weighted by a function of the trial duration.

The analysis of both limit cases suggests that the bias also depends on the trial duration. To confirm this, we extended the trial duration for either limit case. As expected from the analysis, the bias decreases steadily in the simulations (Fig. 5E). We, therefore, predict that the contraction bias can be reduced for sufficiently long trials.

So far, we assumed that the stimulus variance is independent of the stimulus mean. A consequence of this choice is that the bias on either end of the input distribution is largely the same (but with reversed signs). However, behavioral data (see, e.g. Rakitin et al., 1998) shows that the bias increases for stimuli drawn from the upper end of the distribution, a phenomenon usually attributed to *scalar variability*. To capture this in the model, we assume that the stimulus standard deviation linearly increases with the stimulus mean. In these simulations, as expected, the bias increases for a stimulus distribution shifted to higher trial means (Fig. S10).

In summary, we show that the weighted integration of sensory inputs and predictions can be interpreted as a neural manifestation of the contraction bias. While the stimulus and trial-to-trial variability shape the contraction bias, their contributions differ. Moreover, we reveal that the trial duration contributes to the bias.

## Discussion

Our work has been driven by the puzzling question of how the brain integrates top-down feedback predictions with the sensory feedforward bottom-up inputs it constantly receives during behavior. This task may be particularly challenging when the prediction and the sensory information differ (Han and Helmchen, 2023). Conflicting information may be caused by noise in the sensory feedforward inputs or by changes in the environment that could not be predicted. A prominent hypothesis is that how much we rely on our predictions and new sensory evidence is determined by an intricate balance between both, based on how reliable they are (see e.g. Körding and Wolpert, 2004; Yon and Frith, 2021).

This idea is consistent with Bayesian theories on the optimal integration of multiple sensory cues (multisensory integration). Ernst and Banks (2002) showed that to estimate the height of a bar humans combine visual and haptic information in a fashion that minimizes the variance of the final estimate. Similar studies confirmed that animals can optimally combine multiple sensory information by taking into account their uncertainties (Battaglia et al., 2003; Körding and Wolpert, 2004; Alais and Burr, 2004; Rowland et al., 2007; Gu et al., 2008; Fetsch et al., 2012). These behavioral studies were accompanied by neural recordings identifying populations of neurons that can form the basis of multisensory integration (Wallace et al., 1998; Gu et al., 2008; Fetsch et al., 2012).

### Summary of findings

Here, we show that PE neurons can serve as the backbone for estimating the uncertainty of both the feedforward sensory inputs and the feedback predictions (Figs. 2 & 3). In our model, we assume a hierarchy of PE circuits that are feed-forwardly connected through the lower-order memory neuron whose activity encodes the mean of the sensory bottom-up inputs. This local prediction is fed back to the lower-order circuit and at the same time feed-forwarded to the higher-order subnetwork (Fig. 1). With this architecture in place, we show that we rely more strongly on our internal signals when the perceived sensory cues are noisier than the predictions. Moreover, our work suggests that predictions modulate neural activity more at the onset of a new sensory input, even if the stimulus is not noisy. As a consequence, studying neural signatures of predictions in the brain might require experiments that involve sufficiently short trials.

Furthermore, we show that the weighting of sensory inputs and predictions can be biased by neuromodulators, as has been suggested before (see, e.g., Yon and Frith, 2021). In our model, those modulatory signals act through interneurons (Cardin, 2019) whose activities increase in the presence of neuromodulators. When PV neuron activity increases, the network weighs predictions stronger than without modulation. In contrast, when VIP neuron activity increases, the network underestimates the uncertainty of the prediction in a sensory-driven regime, and it underestimates the uncertainty of the sensory input in a prediction-driven regime. Hence, the system leans toward weighting sensory inputs and predictions more equally (Fig. 4A). When SOM and VIP neuron activities are modulated to the same degree, the weighting remains unaffected, suggesting that the individual contributions cancel (Fig. 4B). We show that these findings can be explained by changes in the baseline and gain of PE neurons arising through the modulation of interneuron activity (Fig. 4D). These results can be tested experimentally by optogenetically or pharmacologically stimulating specific interneuron types.

Finally, we illustrate that the weighted integration of feedforward and feedback inputs can be interpreted as a neural manifestation of the contraction bias. We show that the bias is strongly driven by the variances of the sensory cue and the prediction, as well as the trial duration. While the sensory noise increases the contraction bias, the uncertainty in the prediction or long trials decreases the bias (Fig. 5). However, we note that we only consider a neural representation of the stimulus and do not account for other sources of noise, like execution noise, that surely impacts the contraction bias observed in behavioral studies.

### Biological evidence for model choices and assumptions

In our model, we assumed that there are dedicated neurons that encode the variance of the feedforward sensory inputs and the prediction (see also Wilmes et al., 2023). This assumption is consistent with the idea that neurons explicitly encode in their activity the parameters (for instance, mean or variance) of a probability distribution (O’Neill and Schultz, 2010; O’Reilly et al., 2012). However, how variances of signals are represented in the brain is still not comprehensively understood and alternative ideas have been put forward. For instance, it is conceivable that the variance is encoded in a population of neurons, each differently tuned to a specific parameter (Knill and Pouget, 2004). The neurons’ activities represent how close the sensory input is to the preferred (predicted) input of each neuron. Similarly, a neuron’s response variability has been suggested to be related to the uncertainty of sensory stimuli (Hoyer and Hyvärinen, 2002; Ma et al., 2006).

There has been evidence that indeed (population of) neurons can encode uncertainty (Soltani and Izquierdo, 2019). For instance, neurons in the parietal cortex in monkeys encode the degree of confidence in a perceptual decision (Kiani and Shadlen, 2009). Similarly, the firing rate of neurons in the orbitofrontal cortex have been shown to encode confidence irrespective of sensory modality (Masset et al., 2020). Neural signatures of uncertainty have been found in regions of the prefrontal cortex (Rushworth and Behrens, 2008), the rat insular and orbitofrontal cortex (Jo and Jung, 2016), or the dorsal striatum in monkeys (White and Monosov, 2016). Moreover, the accuracy of memory recalls is encoded in single neurons of the human parietal and temporal lobes Rutishauser et al. (2015, 2018).

In our model, we assume a hierarchy of predictions that are locally computed in memory neurons. These memory neurons are consistent with the idea of internal representation neurons hypothesized in predictive processing theories (Bastos et al., 2012; Keller and Mrsic-Flogel, 2018). While it has been hypothesized that these internal representation neurons might be deeper L5 neurons (Bastos et al., 2012; Heindorf and Keller, 2022), there is also evidence that a group of excitatory L2/3 neurons integrates over negative and positive prediction errors (O’Toole et al., 2022).

The core hypothesis of our model is the presence of sensory PE neurons. Those neurons have been found in different cortical areas in various species (Eliades and Wang, 2008; Keller and Hahnloser, 2009; Ayaz et al., 2019; Audette et al., 2021). Moreover, while first only hypothesized theoretically (Rao and Ballard, 1999), the presence of two types of PE neurons, the negative and positive PE neurons, has been confirmed in several recent studies (Keller et al., 2012; Attinger et al., 2017; Jordan and Keller, 2020; Audette et al., 2021). In our model, the nPE neuron inhibits the memory neuron while the pPE neuron excites it, in line with Keller and Mrsic-Flogel (2018). The weights from those PE neurons onto the memory neuron are larger for the lower than the higher PE circuit, so that the lower-order memory neuron evolves faster than the higher-order counterpart. This assumption is consistent with the observation that time constants increase along the cortical hierarchy (Murray et al., 2014; Chaudhuri et al., 2015; Runyan et al., 2017).

If the relation of the paces at which the memory neurons evolve is strongly modulated, the network either shows a bias towards the sensory cues or the prediction (Figs. S5). It has been hypothesized that symptoms in psychiatric diseases may derive from an erroneous uncertainty estimation (Yon and Frith, 2021). For instance, hallucinations may arise from an underestimation of the expectation uncertainty or an overestimation of the sensory uncertainty. Conversely, a fixation on the environment, even when the sensory cues indicate a switch in the environment, may originate from an overestimation of the expectation uncertainty or an underestimation of the sensory uncertainty (Yon and Frith, 2021).

### Neuromodulators and uncertainty

A popular hypothesis is that neuromodulators shape the weighting of sensory inputs and predictions thereof (Yon and Frith, 2021). Theoretical work by Yu and Dayan (2005) suggests that acetylcholine (ACh) correlates with *expected uncertainty*, while noradrenaline (NA) correlates with *unexpected uncertainty*. Expected uncertainty is usually interpreted as known cue-outcome unreliabilities. In contrast, unexpected uncertainty relates to the changes in the environment that produce large PEs outside the expected range of uncertainties (Yu and Dayan, 2005). While in our network the stimulus and trail-to-trial variability can only be loosely interpreted as ‘expected’ and ‘unexpected’ uncertainty, we want to compare the effects of ACh and NA on the weighting with the effects hypothesized in the literature.

It is assumed that NA increases in more volatile environments and enhances bottom-up processes (Hasselmo et al., 1997; Yon and Frith, 2021). In line with this idea, NA blockade impairs cognitive flexibility (Ridley et al., 1981; Janitzky et al., 2015). In recent work by Lawson et al. (2021), it has been shown that humans receiving propranolol (blocking NA) rely more strongly on their expectations and are slower to update these predictions despite new sensory evidence (Yon and Frith, 2021). A main target for noradrenergic inputs is SOM neurons whose activity increases in the presence of NA (reviewed in, e.g., Urban-Ciecko and Barth, 2016; Hattori et al., 2017; Swanson and Maffei, 2019). In our model, activating SOM neurons does not enhance sensory bottom-up input. In a volatile environment, that is, a sensory-driven regime, the system takes into account predictions slightly more than without SOM modulation (Figs. 4A and S6).

However, we note that in our simulations, we assumed that neuromodulators act globally, that is, on the interneurons in both the lower and the higher PE circuit. While this agrees with the view that neuromodulators can control network states globally, there is also evidence that they can have a more local, finely adjusted impact on neural circuits (Nadim and Bucher, 2014). In our model, increasing SOM activity only in the lower-order circuit shows a slight enhancement of the sensory weight (Fig. S7), that is, the bottom-up inputs. This suggests that whether a neuromodulator biases the network toward feedforward bottom-up or feedback top-down inputs depends on its spatial and temporal scale of influence.

Similarly to NA, ACh has also been shown to enhance bottom-up, feedforward inputs (reviewed in, e.g., Yu and Dayan, 2005; Marshall et al., 2016). For instance, subjects relied more strongly on prior beliefs when given cholinergic receptor antagonists (Marshall et al., 2016). A major target for cholinergic inputs is VIP neurons whose activity increases in the presence of ACh (reviewed in, e.g., Wester and McBain, 2014; Hattori et al., 2017; Swanson and Maffei, 2019). In our model, activating VIP neurons globally only enhances bottom-up input in stable environments for noisy stimuli. However, increasing VIP activity only in the higher-order PE circuit generally enhances sensory bottom-up inputs (Fig. S7).

### Limitations & future steps

As for any computational model, we brush over several biological details to keep the model simple and interpretable. However, those details, while beyond the scope of this study, may be well investigated in future work. For instance, neuromodulatory systems have been suggested to gate plasticity (Pawlak et al., 2010). In a recent study by Jordan and Keller (2023), locus coeruleus (LC) axon activity is shown to correlate with the magnitude of unsigned visuomotor prediction errors. The authors hypothesize that LC output modulates the learning rate at which the internal model evolves (Jordan and Keller, 2023). In our model, we do not consider precision-weighted PEs (but see Wilmes et al., 2023; Granier et al., 2023). Hence, a sensible extension to our work would be to adjust the weights from PE neurons onto the memory neurons by a function of the stimulus and prediction uncertainty, respectively. This would allow us to compare our results more closely to work showing that ACh and NA can adjust the rate at which new sensory evidence is incorporated when environments change (Marshall et al., 2016; Bruckner et al., 2022).

Our model suggests *one* potential neuronal circuit mechanism for the estimation of sensory inputs and predictions. However, in the light of evidence showing that the integration of feedforward and feedback inputs is species- and modality-dependent, it is conceivable that a plethora of neural mechanisms are used in neural circuits. For multisensory integration, Wong et al. (2023) showed that in Drosophila larva the chosen cue-combination strategy varies depending on the type of sensory information available. Also, humans put typically more weight on visual than auditory cues (Battaglia et al., 2003; Alais and Burr, 2004), but trust vestibular information more than visual information about head direction (Butler et al., 2010), a finding also observed for monkeys (Fetsch et al., 2009). Moreover, Summerfield et al. (2011) showed that humans diverge from an optimal Bayesian strategy in very volatile environments and act according to their experience in the last trial. It has been suggested that the brain may use different strategies to combine signals depending on the task demands (O’Reilly et al., 2012). While these behavioral results cannot speak to the underlying circuit mechanisms, it is conceivable that neural implementations for the integration of feedforward and feedback inputs may also vary.

Furthermore, while we provide a neuronal circuit model for estimating the mean and variance of both sensory signals and predictions, we do not explicitly model the weighting of inputs. A respective neural circuit model would require nested inhibitory interneurons providing divisive inhibition. How this subnetwork interacts with the PE circuits, and how the presence of neuromodulators acting on the interneurons directly involved in the weighting, impacts our findings is subject to future work.

### Relation to other work & conclusions

Many normative models have been proposed for state estimation and prediction under uncertainty (Soltani and Izquierdo, 2019), ranging from the classical Kalman filter to more recent models like *Bayes Factor Surprise* (Liakoni et al., 2021). For instance, the Bayes factor surprise formularizes the trade-off between integrating new observations in an existing belief system and resetting this belief system with novel evidence. The surprise factor captures how much an animal’s current belief deviates from the new observation.

In recent years, normative models have been squared with biological constraints. For instance, Kutschireiter et al. (2023) showed that a *Bayesian ring attractor* model can encode uncertainty in the amplitude of the network activity and matches the performance of a circular Kalman filter when the recurrent connections are tuned appropriately. In other seminal work, it has been proposed that Bayesian inference in time can be linked to the dynamics of leaky integrate-and-fire neurons with spike-dependent adaptation (Deneve, 2008).

Here, we proposed an alternative view in which PE neurons serve as the backbone for estimating both the uncertainty of the feedforward sensory stimuli arising from the external world and the feedback signals carrying predictions about the same feedforward inputs our brains receive. Our work is an important step toward a better understanding of the brain’s ability to integrate these unreliable feedforward and feedback signals that often do not match perfectly.

## Models and methods

### Network model

The mean-field network model consists of a *lower* and *higher* PE circuit (Fig. 1C-D). Each PE circuit contains an excitatory nPE neuron and pPE neuron (N_nPE_ = N_pPE_ = 1), as well as inhibitory neurons. The inhibitory neurons comprise PV, SOM and VIP neurons (N_SOM_ = N_VIP_ = 1, N_PV_ = 2), further explained in Hertäg and Clopath (2022). In addition to the core PE circuit, each subnetwork also includes one memory neuron *M* and one variance neuron *V* .

The excitatory neurons in the PE circuit are simulated as two coupled point compartments, representing the soma and the dendrites of elongated pyramidal cells. All other neurons are modeled as point neurons. The activities of all neurons are represented by a set of differential equations describing the network dynamics.

The dynamics of the neurons in the lower and higher PE circuits (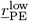 and 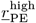) are given by

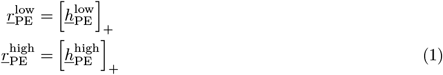

with

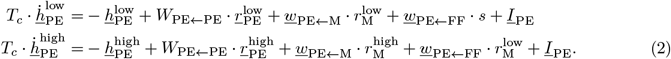

We follow the notation that column and row vectors are indicated by letters with an underscore <u>•</u>, matrices are denoted by capital letters, and scalars are given by small letters without an underscore. Furthermore, a time derivative (e.g., 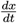) is denoted by a dot above the letter (e.g., 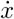). The rate vector 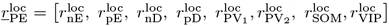 with loc ∈ [low, high] contains the activities of all neurons or compartments in the PE circuit (soma of nPE/pPE neurons: nE/pE, dendrites of nPE/pPE neurons: nD/pD). The network receives time-dependent stimuli *s* and neuron/compartment-specific external background input *<u>I</u>*_PE_. The connection strengths between the *pre*-synaptic population and the neurons of the PE circuit are denoted by *W*_PE←pre_ (if *r* is a vector) or *<u>w</u>*_PE←*pre*_ (if *r* is a scalar). The activities of the neurons evolve with time constants summarized in *T*_*c*_.

The activities of the lower and higher memory (M) neuron evolve according to a perfect integrator. The M neurons receive synapses from both nPE and pPE neurons of the same subnetwork,

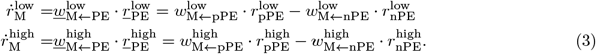

The activities of the lower and higher V neuron evolve according to a leaky integrator with quadratic activation function. The variance neurons receive synapses from both nPE and pPE neurons of the same subnetwork,

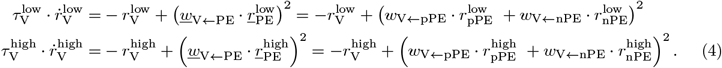

All values for neuron and network parameters, details on the model equations for the mean-field and the population network, as well as supporting analyses can be found in the supplementary material.

### Weighting of sensory inputs and predictions

We arithmetically calculated the weighted output of sensory inputs and predictions, *r*_out_, based on ideas of Bayesian multisensory integration (see, e.g. Pouget et al., 2013),

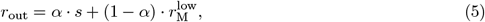

where *α* denotes the sensory weight (that is, the reliability of the sensory input) and is given by

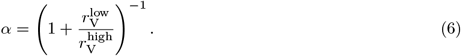

### Inputs

The network receives feedforward stimuli *s* that may vary between trials. To account for noise, each stimulus is composed of N_in_ constant values drawn from a normal distribution with mean *μ*_in_ and standard deviation *σ*_in_, and are presented for N_step_ consecutive time steps. To account for changes in the environment, *μ*_in_ is drawn from a uniform distribution *U* (*a, b*) with mean 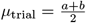 and standard deviation 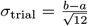 . The parameterization of both distributions varies across the experiments. All stimulus/input parameters can be found in the supplementary material.

### Simulations

All simulations were performed in customized Python code written by LH. Source code to reproduce the simulations, analyses, and figures will be available after publication at https://github.com/lhertaeg/weighted_sensory_prediction. Differential equations were numerically integrated using a 2^nd^-order Runge-Kutta method. Neurons were initialized with *r* = 0*/s*. Further details and values for simulation parameters can be found in the supplementary material.

## Acknowledgments

We thank Inês C. Guerreiro for comments on earlier versions of this manuscript. This work was supported by Deutsche Forschungsgemeinschaft (DFG) Grant 460088091, Biotechnology and Biological Sciences Research Council (BBSRC) Grants BB/N013956/1 and BB/N019008/1, Wellcome Trust Grant 200790/Z/16/Z, Simons Foundation Grant 564408, and Engineering and Physical Sciences Research Council (EPSRC) Grant EP/R035806/1.

## Supporting Information

### A Detailed Methods

In the following, we describe in more detail the equations for the dynamics of the neurons in the prediction-error circuit, as well as the memory and variance neurons. We then provide the connectivity of the network and the inputs to the neurons for both the mean-field and multi-cell population model. Finally, to ensure reproducibility, we summarize all simulation parameters used for the results shown in the figures.

#### A.1 Network model

The network model consists of a *lower* and *higher* mean-field PE circuit (Fig. 1). Each PE circuit contains an excitatory nPE neuron and pPE neuron (N_nPE_ = N_pPE_ = 1), as well as inhibitory neurons. The inhibitory neurons comprise PV, SOM and VIP neurons (N_SOM_ = N_VIP_ = 1, N_PV_ = 2). As has been shown in Hertäg and Clopath (2022), we need two soma-targeting interneurons, one receiving the sensory input and one receiving the prediction, to obtain a perfect nPE and pPE neuron in the same recurrent network. We, therefore, used two PV neurons (as suggested in the original paper). In addition to the core PE circuit, each subnetwork also includes one memory neuron *M* and one variance neuron *V* .

In Figure 2 and the corresponding supporting figures, only the lower subnetwork is simulated. In Fig. S2, we replaced this lower mean-field PE circuit with a heterogeneous multi-cell population model containing 200 neurons (N_SOM_ = N_VIP_ = N_PV_ = 20, 140 excitatory neurons). In Fig. S3, the lower PE circuit comprises 1000 copies of the mean-field network to account for selectivity.

In the following, we describe the dynamics of the neurons/compartments in the mean-field network. The equations for the population PE circuit (Fig. S2) are directly deduced from the mean-field equations and can also be found in (Hertäg and Clopath, 2022).

##### A.1.1 Prediction-error network model

Each excitatory pyramidal cell (that is, nPE or pPE neuron) is divided into two coupled compartments, representing the soma and the dendrites, respectively. The dynamics of the firing rates of the somatic compartments *r*_nE_ (nPE neuron) and *r*_pE_ (pPE neuron) obey (Wilson and Cowan, 1972)

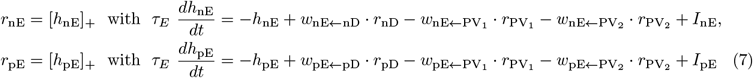

where *τ*_E_ denotes the excitatory rate time constant (*τ*_E_=60 ms), the weights *w*_nE←nD_ and *w*_pE←pD_ describe the connection strength between the dendritic compartment and the soma of the same neuron, and 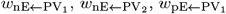 and 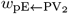 denote the strength of somatic inhibition from PV neurons. The overall input *I*_nE_ and *I*_pE_ comprise the external background and feedforward inputs (see “Inputs” below). Firing rates are rectified to ensure positivity ([*•*]_+_).

The dynamics of the activity of the dendritic compartments *r*_nD_ (nPE neuron) and *r*_pD_ (pPE neuron) obey (Wilson and Cowan, 1972)

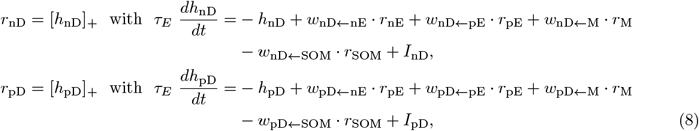

where the weights *w*_nD←nE_, *w*_nD←pE_, *w*_pD←nE_ and *w*_pD←pE_ denote the recurrent excitatory connections between PCs. *w*_nD←SOM_ and *w*_pD←SOM_ represent the strength of dendritic inhibition from the SOM neuron. *w*_nD←M_ and *w*_pD←M_ denote the strength of connection between the memory neuron and the dendrites. The overall inputs *I*_nD_ and *I*_pD_ comprise fixed, external background inputs (see “Inputs” below). We assume that any excess of inhibition in a dendrite does not affect the soma, that is, the dendritic compartment is rectified at zero.

Similarly, the firing rate dynamics of each interneuron is modeled by a rectified, linear differential equation,

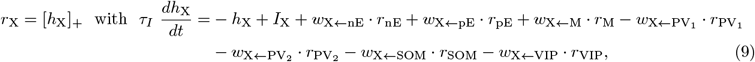

where *r*_X_ denotes the firing rate of interneuron type *X*, and the weight *w*_X←Y_ denotes the strength of connection between the presynaptic neuron *Y* and the postsynaptic neuron *X* (*X* ∈ *{*PV_1_, PV_2_, SOM, VIP*}, Y* ∈ *{*nPE, pPE, PV_1_, PV_2_, SOM, VIP, M*}*). The rate time constant *τ*_*I*_ was chosen to resemble a fast GABA_A_ time constant, and set to 2 ms for all interneuron types included. The overall input *I*_X_ comprises fixed, external background inputs and feedforward sensory inputs (see “Inputs” below).

##### A.1.2 Memory and variance neuron

In addition to the core PE circuit, we simulate a memory neuron *M* and a variance neuron *V*. The memory neuron is modeled as a perfect integrator, receiving synapses from both the nPE and pPE neuron,

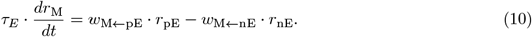

*w*_M←pE_ denotes the connection strength between the pPE neuron and the memory neuron, and *w*_M←nE_ denotes the connection strength between the nPE neuron and the memory neuron. The time constant *τ*_*E*_ = 60 ms.

The dynamics of the variance neuron obeys a non-linear differential equation with leak term,

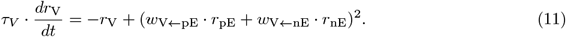

The weight *w*_V←pE_ represents the connection strength between the pPE neuron and the variance neuron, while *w*_V←nE_ denotes the connection strength between the nPE neuron and the variance neuron. To ensure that the V neuron encodes the variance, we chose a quadratic activation function. In Fig. S5, we used a linear activation function to investigate the impact of the input-output transfer function on the weighting of sensory inputs and predictions. The time constant *τ*_*V*_ was 5 *s* in the mean-field model, 2 *s* in the heterogeneous multi-cell population model (Fig. S2), and 0.5 *s* in the network model with selectivity (Fig. S3).

##### A.1.3 Weighted output

The weighted output *r*_out_(*t*) is a linear combination of the current sensory input *s*(*t*) and the activity of the memory neuron, *r*_M_(*t*), inspired by Bayesian multisensory integration (see, e.g. Pouget et al., 2013),

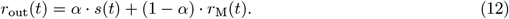

How strongly either the sensory input or the prediction thereof contributes to the output is denoted by the sensory weight *α*,

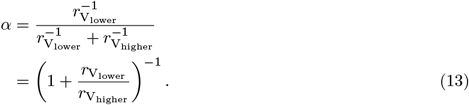

#### A.2 Connectivity

##### A.2.1 Connections between neurons of the PE circuit

The connectivity between neurons of the PE circuit, both for the mean-field and the population network, were taken from (Hertäg and Clopath, 2022). We considered three mean-field networks (see Table 2) that differed in terms of the inputs (feedforward vs. feedback) onto the SOM and VIP neurons, and, hence, in their connectivity that established an E/I balance in the excitatory neurons.

##### A.2.2 Connections between the PE circuit and the M neuron

While the nPE neurons inhibit the M neuron, the pPE neurons excite it. To ensure that the activities of the memory neurons represent the mean of the sensory stimuli in the lower PE circuit and the mean of the prediction in the higher subnetwork, respectively, the net effect of nPE and pPE neurons must cancel in the steady state (see Analysis in B.2). Hence, the weights need to account for the neurons’ potentially different gain factors (*g*_nPE_ and *g*_pPE_) and the neuron numbers (*N*_nPE_ and *N*_pPE_):

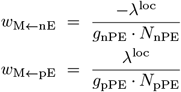

where *λ*^loc^ denotes a weight for the lower or higher-order PE circuit, loc ∈ *{*lower, higher*}*. In the lower PE circuit, *λ*^lower^ = 3 · 10^−3^ for the mean-field model in Fig. 2 and *λ*^lower^ = 4.5 · 10^−2^ for Figs. 3-5. In the higher PE circuit, *λ*^higher^ = 7 · 10^−4^.

For the mean-field networks (*N*_nPE_ = *N*_pPE_ = 1), the gain factors *g*_nPE_ and *g*_pPE_ are given in Table 1. For the population network, the gain factors for all PE neurons are shown in Fig. S0.

**Table 1.** Gain factors for nPE and pPE neurons in three different mean-field networks (MFN). Each MFN differs with respect to the inputs onto SOM and VIP neurons. The interneurons either receive the feedforward (FF) or feedback (FB) input. All numbers are rounded to the first digit.

| Network | MFN 1 | MFN 2 | MFN 3 |
| --- | --- | --- | --- |
| | FF → SOM<br>FB → VIP | FB → SOM<br>FF → VIP | FF → SOM<br>FF → VIP |
| nPE | 1 | 1.7 | 2.5 |
| pPE | 1 | 1.7 | 2.5 |

**Table 2.** *w*_X←M_ for the post-synaptic SOM and VIP neurons in all three mean-field networks considered.

| Network | MFN 1 | MFN 2 | MFN 3 |
| --- | --- | --- | --- |
| | FF → SOM<br>FB → VIP | FB → SOM<br>FF → VIP | FF → SOM<br>FF → VIP |
| SOM | 0 | 1 | 0 |
| VIP | 1 | 0 | 0 |

**Figure S0.**
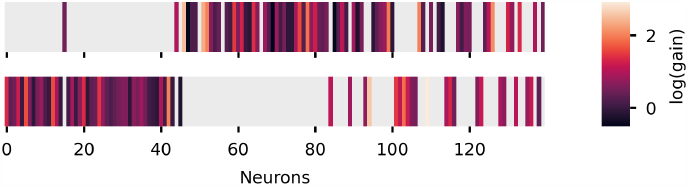
Gain factors of nPE and pPE neurons in the multi-cell population model. The logarithm of the gain factors of nPE (top) and pPE (bottom) neurons in the multi-cell population model from (Hertäg and Clopath, 2022). The network contains 67 nPE neurons and 66 pPE neurons. The remaining excitatory neurons were not classified as PE neurons and were not connected to the *M* neuron.

The memory neuron *M* connects to the post-synaptic neurons *X* in the PE circuit with the connection strength *w*_X←M_ = 1, if a connection exists, *w*_X←M_ = 0 otherwise. In all mean-field networks and the population network, the dendrites of nPE and pPE neurons and one of the two (populations of) PV neurons receive connections from the memory neuron. Furthermore, we assume that the *M* neuron does not excite the soma of PCs. Whether the SOM or VIP neurons are the target of the feedback projections depend on the specific mean-field network (see Table 2). In the multi-cell population model, 30% of the SOM neurons and 70% of the VIP neurons receive input from the memory neuron.

##### A.2.3 Connections between the PE circuit and the V neuron

Both nPE and pPE neurons excite the *V* neuron. To ensure that the activity of the V neuron represents the variance of the input (see Analysis in B.2), the weights must account for differences in the gains (*g*_nPE_ and *g*_pPE_, see Table 1 and Fig. S0) and numbers (*N*_nPE_ and *N*_pPE_) of the PE neurons,

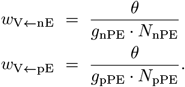

The factor *θ* denotes the unscaled weight, and can be chosen to compensate for potential deviations from the definition of the variance as a result of a quadratic activation function. In the mean-field network, *θ* = 1 because nPE and pPE neuron activity is mutually exclusive, and, hence, the cross-term nPE· pPE would be zero (under the assumption that they have a negligible baseline activity). This is also true for the multi-cell population model. Each PE neuron receives the same feedforward stimulus and contributes only a small (scaled) fraction to the overall PE (adding up to the same PE used in the mean-field network).

However, in Supp Fig. S3, each mean-field network receives a different stimulus *s*_*i*_ drawn from a distribution at time *t*. While the stimuli below the mean of the distribution activate the nPE neurons (each located in a different mean-field network), the stimuli above the mean of the distribution activate the pPE neurons (each located in a different mean-field network). Because the V neuron first sums all the contributions from the PE neurons before applying the non-linearity, its steady state activity is similar to the squared sum of the *averaged* nPE and *averaged* pPE neuron activity. In this case, *θ* must be chosen such that deviations from the true variance can be mitigated or fully corrected. The true *θ* depends on the distribution at hand. In our simulations, we used a uniform distribution *U* (*a, b*), in which case *θ* can be derived from

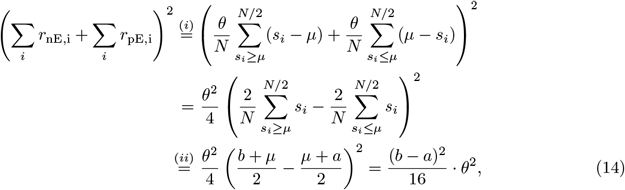

where we assumed that (i) the number of nPE and pPE neurons is equal, and (ii) this number goes to infinity. Comparing eq. (14) with the equation for the variance of a uniform distribution, 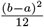, we get 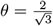 .

#### A.3 Inputs

Each neuron (type) receives an overall input *I*_*i*_,

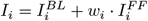

where 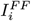 denotes a feedforward input and 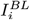 represents an external background input that ensures reasonable baseline firing rates in the absence of sensory inputs and predictions thereof. In the case of the mean-field network, these inputs were set such that the baseline firing rates are *r*_pE_ = *r*_pD_ = *r*_nE_ = *r*_nD_ = 0 *s*^−1^ and *r*_P_ = *r*_S_ = *r*_V_ = 4 *s*^−1^. In the case of the population network, we set the external inputs of all neuron types to 5 *s*^−1^, while the background inputs to the dendrites were computed such that the dendrites are inactive during baseline.

The feedforward input is either the direct sensory input *s* for the lower PE circuit, or the activity of the M neuron, *r*_M_, for the higher PE circuit. In general, for the three mean-field networks tested, we chose *w*_*i*_ = 1− *w*_X←M_ (see Table 2). In the population network, 70% of the SOM neurons and 30% of the VIP neurons receive the feedforward input.

#### A.4 Simulations

All simulations were performed in customized Python code written by LH. Source code to reproduce the simulations, analyses and figures will be available after publication at https://github.com/lhertaeg/weighted_sensory_prediction. Differential equations were numerically integrated using a 2^nd^-order Runge-Kutta method. Neurons were initialized with *r* = 0*/s*.

The qualitative results were fairly robust to the choice of the simulation parameters and are here stated merely to ensure the reproducibility of all figures. However, we note that we made use of PE circuits that had been trained on steady state inputs (Hertäg and Clopath, 2022). Hence, we must simulate the network long enough to ensure that the PE neurons reach their steady state. Moreover, the lower-level M neuron must evolve faster than the higher-level M neuron as indicated in Fig. S5. Finally, the time constant of the V neurons must be of the same magnitude as the trial duration.

In the following, we give all figure-specific parameters not directly visible or mentioned in the figures and captions. Furthermore, to increase readability, we do not include units for the parameters. All units can be deduced from the equations above. We simulated the network in Figures 1 and 2 for 10^5^ simulation time steps. In Figure 2, we presented 200 constant values, each 500 time steps long. In Figure 3 (including supporting figures) we simulated 100 trials, while in Figures 4 to 5 (including supporting figures), we simulated 200 trials, each 5000 time steps long. In a trail, 10 constant values were drawn from a normal distribution 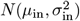, each 500 time steps long. The stimulus mean was drawn from an uniform distribution, *U* (*a, b*), with mean *μ*_in_ and variance 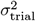 .

**Figure 1:** Constant prediction (fixed) = 5, input mean ∈ [0, 10], input standard deviation 0.

**Figure 2:** Inputs drawn from a uniform distribution. B and C: input standard deviation fixed at 4.5 when mean is varied, input mean fixed at 4.5 when input variance is varied.

**Figure 2 Supplementary Fig. 1:** Inputs drawn from different distributions with mean of 5 and variance of 4. Number of repetitions with different seeds: 20.

**Figure 2 Supplementary Fig. 2:** Inputs drawn from a uniform distribution with mean of 5 and variance of 4. Time step was 0.1. The connections from the PE neurons to the M or V neuron were altered by a factor *γ* drawn from a normal distribution. If not stated otherwise, the mean of this normal distribution was 1 and the variance 0, while the connection probability was 1. Number of repetitions with different seeds: 10.

**Figure 2 Supplementary Fig. 3:** Number of time steps were 4000. Number of identical mean-field networks: 1000.

**Figure 3:** 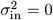 and *U*(1, 9), 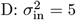 and *U*(5, 5), F: 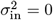 and *U*(*a, b*) was parameterized such that the trail variability was 3.

**Figure 3 Supplementary Fig. 1:** Switch of input statistics occurs after 50 trails. State 1: stimulus ∈ *N* (*μ*_in_, 0) with *μ*_in_ ∈ *U* (5, 5), State 2: stimulus ∈ *N* (*μ*_in_, 3) with *μ*_in_ ∈ *U* (5, 5), State 3: stimulus ∈ *N* (*μ*_in_, 3) with *μ*_in_ ∈ *U* (0, 10), State 4: stimulus ∈ *N* (*μ*_in_, 0) with *μ*_in_ ∈ *U* (0, 10).

**Figure 3 Supplementary Fig. 2:** *μ*_in_ = 5 and 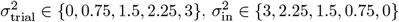. A: scaling factors of *w*_M←PE_ were 0.3 and 7.

**Figure 4:** *μ*_in_ = 5. A, top: 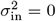 and 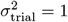. A, bottom: 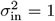 and 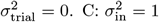 and 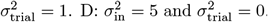. Additional input (perturbation) was either fixed at 0.5 (A, D) or systematically varied between − 1 and 1 (C), and was *on* for the last 50% of the trials. To estimate changes in baseline and gain of nPE and pPE neurons, we fitted a linear function to the PE neuron activity for the input range [0, 2.5].

**Figure 4 Supplementary Fig. 1:** All parameters are from Figure 4 A.

**Figure 4 Supplementary Fig. 2:** In the default setting baseline was 0 and gain of PE neurons was 1.

The results (left) were computed for baselines ∈ [0, 3], while the results (right) were computed for gains [0.5, 1.5]. The results are based on the Eqs. (22), (25), (27) and (30).

**Figure 4 Supplementary Fig. 3:** All parameters are from Figure 4 D.

**Figure 5:** A: *σ*_in_ ∈ *{*1, 7*}, μ*_in_ ∈ *U* (15, 25). B: *σ*_in_ ∈ [0, 8], *μ*_in_ ∈ *U* (15, 25), and *σ*_in_ = 5, *μ*_in_ ∈ *U* (15, *b*) with *b* ∈ [20, 48]. C: *σ*_in_ ∈ {2, 5}, *μ*_in_ ∈*U* (15, 15). D: *σ*_in_ = 0, *μ*_in_ ∈*U* (15, 25) or *U* (10, 30). E: *σ*_in_ = 0, *μ*_in_ ∈*U* (15, 25), or *σ*_in_ = 5, *μ*_in_ ∈*U* (15, 15), Time steps per trail increased from 5000 to 10^4^.

**Figure 5 Supplementary Fig. 1:** We used two different uniform distributions *U* (15, 25) and *U* (25, 35), and introduced scalar variability so that *σ*_in_ is a linear function of *μ*_in_. Specifically, we chose *σ*_in_ = [*μ*_in_ − 14]_+_.

## B Supporting analyses

We first describe a simplified model and show that the M neuron represents the mean, while the V neuron represents the variance of the feedforward input. We then investigate the impact of the gain and baseline of PE neurons on estimating the mean and variance. Furthermore, we use the simplified model to discuss the effect of neuromodulators in our network. Finally, we reveal the connection between the sensory weight and the contraction bias.

### B.1 Activity of M and V neuron in a simplified model

To show that the M neuron encodes the mean, while the V neuron encodes the variance of the feedforward input, we resume a toy model in which the activity of the nPE and pPE neuron is replaced by its ideal output

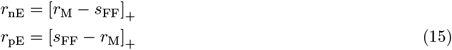

with *s*_FF_ denoting the time-dependent feedforward input. The activity of the M neuron can then be described as

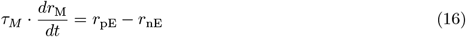

If *r*_M_ ≥ *s*_FF_, we get

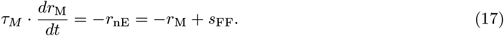

If *r*_M_ ≤ *s*_FF_, we also get

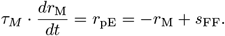

Hence, the activity of *r*_M_ is given by

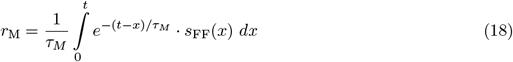

for zero activity at time *t* = 0. In the limit of *t* → ∞ (steady state), this is the exponential moving average of the feedforward input, *E*(*s*_FF_).

With the simplified activity of the nPE and pPE neuron, the activity of the V neuron can then be described as

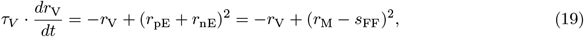

leading to the time-dependent solution

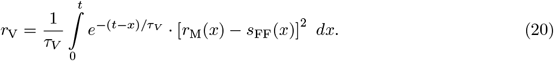

In the limit of *t* → ∞, *r*_V_ approaches *E*(*s*_FF_ − *E*(*s*_FF_)).

### B.2 Impact of PE neurons’ gain on estimating mean and variance

The gains of the PE neurons, if not equal between the nPE and pPE neuron on average, can bias the activity of both the M and V neuron. To show this, we resume the toy model from section B.1.

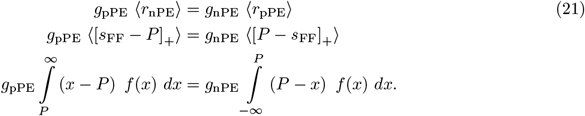

Here, *P* denotes the prediction encoded in the M neuron, and *f* (*x*) is the distribution of feedforward inputs. In case of a uniform distribution, *f* (*x*) = 1*/*(*b* − *a*) for *x* ∈ [*a, b*] and 0 otherwise, we get

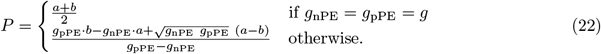

Hence, the mean of the feedforward input is overpredicted when *g*_nPE_ *< g*_pPE_. Similarly, the mean of the feedforward input is underpredicted when *g*_nPE_ *> g*_pPE_ (Fig. S7).

Likewise, the variance is affected by the gain of the nPE and pPE neuron,

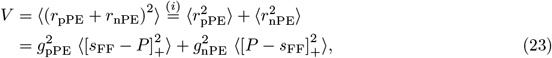

where we assume (i) that both the nPE and pPE neuron have a zero baseline activity. In case of a uniform distribution, we get

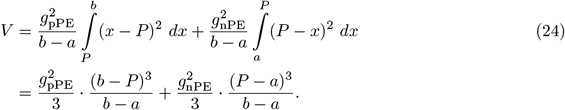

Inserting eqs. (22) yields

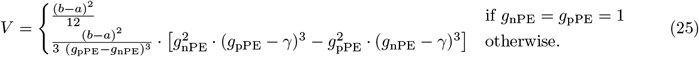

with 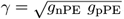. Hence, the variance of the feedforward input is overpredicted when *g*_nPE_ *>* 1 or *g*_pPE_ *>* 1. Similarly, the variance of the feedforward input is underpredicted when *g*_nPE_ *<* 1 or *g*_pPE_ *<* 1 (Fig. S7).

### B.3 Impact of PE neurons’ baseline on estimating mean and variance

The baselines of the PE neurons, if not equal between the nPE and pPE neuron on average, can bias the activity of both the M and V neuron. By means of the toy model from section B.1, we can write

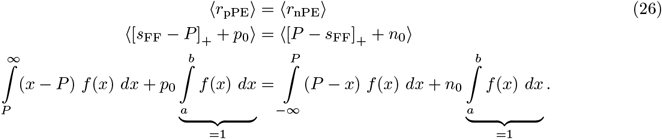

*n*_0_ and *p*_0_ denote the baseline activity of the nPE and pPE neuron, respectively. In case of a uniform distribution (c.f. B.2), we get

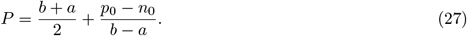

Thus, the M neuron encodes the true mean of the feedforward input only if *p*_0_ = *n*_0_. As a result, the mean is overpredicted if *p*_0_ *> n*_0_. Likewise, the mean is underpredicted if *p*_0_ *< n*_0_ (see Fig. S7).

With non-zero baseline activities, the steady state activity of the V neuron is given by

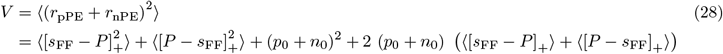

In case of a uniform distribution *U* (*a, b*), this expression yields

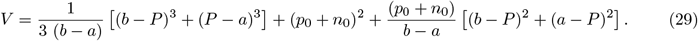

Inserting the expression for P (Eq. 27) which is itself a function of the baseline activities, gives

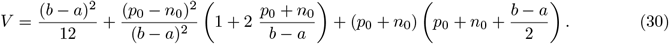

Thus, for the V neuron to encode the variance unbiased, *n*_0_ = *p*_0_ = 0. The variance is overpredicted if either *n*_0_ *>* 0 or *p*_0_ *>* 0 (see Fig. S7).

### B.4 Modelling the impact of neuromodulators on the sensory weight

We modeled the presence of a neuromodulator by simulating an additive excitatory input onto (groups of) interneurons. These interneurons, in turn, modulate the gain and baseline of PE neurons. As shown in sections B.2 and B.3, changes in the input-output transfer function of the PE neurons may bias the variance estimation in the network, and, hence, the sensory weight. Thus, understanding changes in the sensory weight requires an understanding of whether and how different types of interneurons change the PE neurons.

If a neuromodulator only acts on interneurons of the lower-level subnetwork, the sensory weight changes as a consequence of the modulated firing rates of the lower-level and higher-level *V* neurons. The lower-level *V* neuron is directly affected by the changes in the lower-level PE neurons and indirectly affected by changes in the *M* neuron of the same network. The higher-level *V* neuron is also affected by a neuromodulator acting in the lower-level subnetwork because the lower-level *M* neuron projects onto the neurons in the higher-level PE circuit. Hence, if the lower-level *M* neuron represents a biased mean *μ ± δμ*, the variance estimation will be biased as well. This can be seen directly from the definition of the variance,

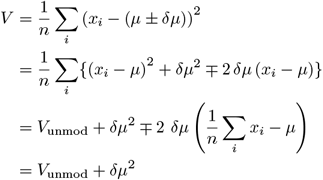

In contrast, if a neuromodulator only acts on interneurons of the higher-level subnetwork, the sensory weight changes as a consequence of the modulated firing rates of the higher-level *V* neuron. The higher-level *V* neuron is directly affected by the changes in the higher-level PE neurons and indirectly affected by changes in the *M* neuron of the same network.

Together, this suggests that whether the sensory weight decreases, increases, or remains the same in the presence of a neuromodulator depends on several factors:

- Does the neuromodulator act on the lower-level or higher-level subnetwork (that is, local impact), or does the neuromodulator act on both to the same degree (that is, global impact)?
- Which interneuron type/s is/are affected by the neuromodulator? And are these interneurons inhibited or excited by the neuromodulator?
- How are these interneurons embedded in the network, that is, what are the connectivity and the inputs to those neurons?

As a result, different neuromodulators may have the same effect on the sensory weight or the same neuromodulator may have different effects depending on brain area, species, etc..

### B.5 Sensory weight and contraction bias

In the simulations, we define the bias as the trial-averaged difference between the weighted output and the true stimulus. For the sake of simplicity, we use *r*_out_ at the end of a trial, *T*, as a proxy for the trial average in the subsequent analysis. Hence,

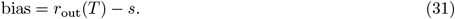

To investigate how the bias depends on the sensory weight and potentially other factors, let us resume a toy model in which we assume that the prediction decays exponentially with time constant *τ* to a presented constant stimulus value, *s*,

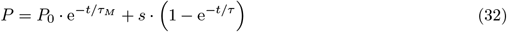

with *P*_0_ describing the initial value at time *t* = 0. Let us further assume that within a trial with trial duration *T*, the stimulus value changes *n* times (*T* = *n* · ∆*t*). The prediction during the presentation of the *n*th stimulus value can be expressed as

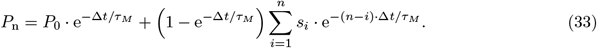

To obtain an estimate for the prediction at the end of a trial, *P*_n_ must be averaged over the stimulus distribution, *⟨P*_n_*⟩*_*s*_. For the sake of simplicity, let us assume the stimulus values are drawn from a uniform distribution 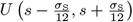 . Moreover, we assume that the initial state, *P*_0_, at the beginning of a new trial is drawn from a uniform distribution 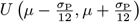 . With these assumptions, *⟨P*_n_ *⟩*_*s*_ is given by

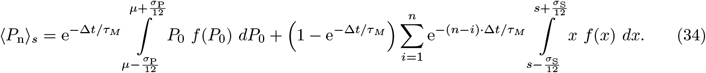

Solving the integrals yield

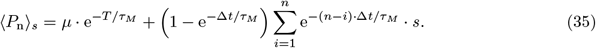

Making use of the geometric series, the expression simplifies to

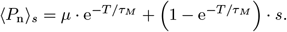

Inserting the expression in the equation for the weighted output yields

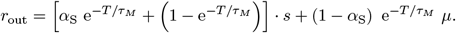

Hence, the bias in our toy model can be expressed by

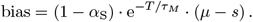

The absolute slope 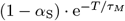indicates how strong the bias is. It depends on the sensory weight *α*_S_, the trial duration *T* and time constant *τ*_*M*_ . Please note that the sensory weight is a function of the trial duration itself (see Fig. 3F). However, for illustration purposes, we take *α*_S_ to be constant.

In this toy model, if the variance of the prediction is zero (that is, in a prediction-driven input regime), *α*_S_≈ 0, and, consequently, the bias is 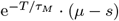 Thus, the bias is independent of the stimulus variance (see Fig. 5 D).

Likewise, if the variance of the sensory stimulus is zero (that is, in a stimulus-driven input regime), *α*_S_ ≈1, and, consequently, the bias approaches 0 if the neurons reach their steady state. Thus, decreasing or increasing the trail variance does not have an effect on the bias (see Fig. 5 C).

## C Supplementary Figures

**Figure S1.**
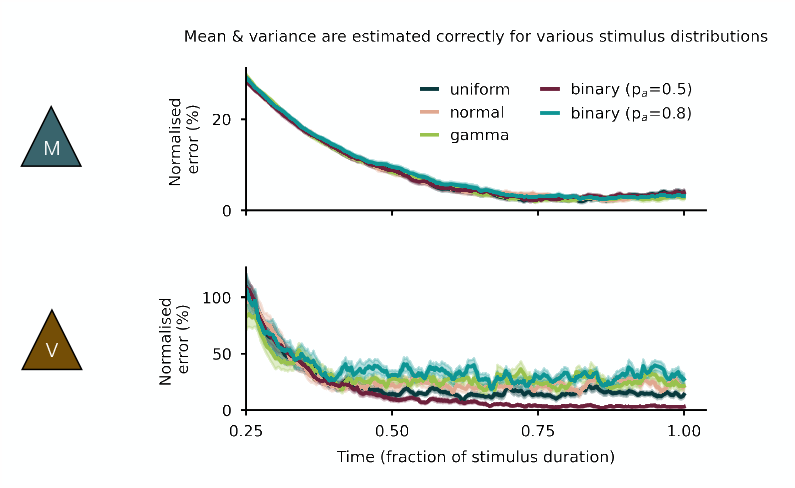
Estimating mean and variance of different stimulus distributions. Top: The normalised absolute difference between the averaged mean and the activity of the M neuron decreases to a near-zero level for all stimulus distributions tested. Bottom: The normalised absolute difference between the averaged variance and the activity of the V neuron decreases with small differences between the distributions tested. Parametrisation of the uniform distribution as in Fig. 2.

**Figure S2.**
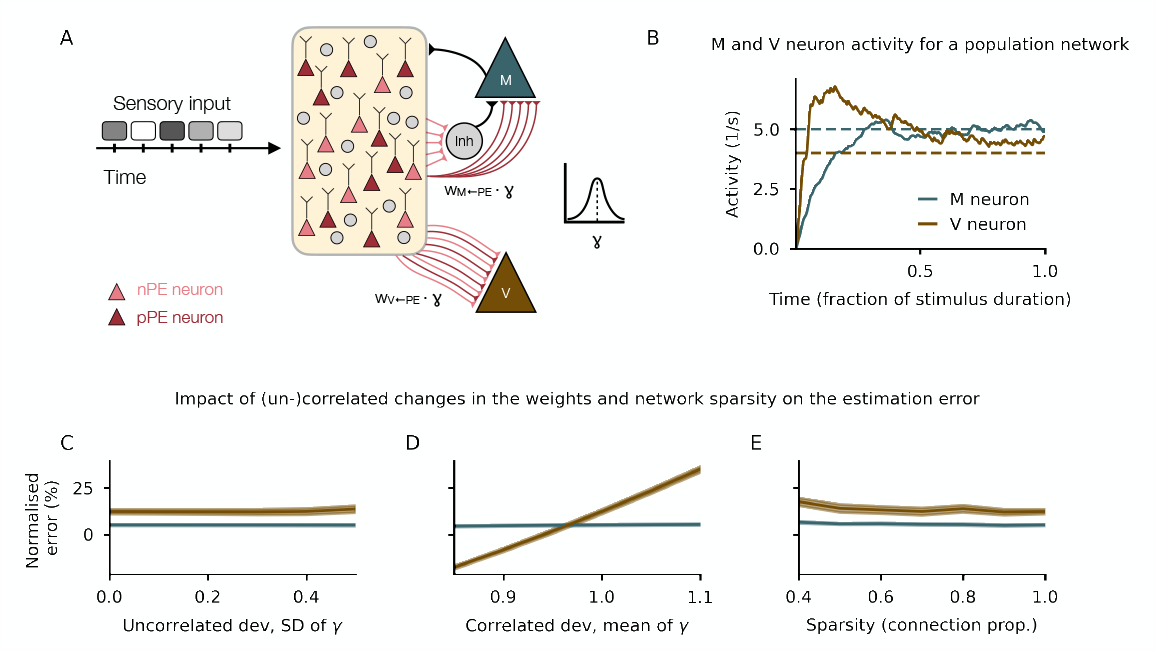
Estimating mean and variance of sensory stimuli in a rate-based multi-cell population network. **(A)** Illustration of the rate-based multi-cell population network and the stimuli over time. The weights from the PE neurons onto the M or V neuron are scaled by a factor *γ* drawn from a normal distribution *N* (*μ*_*γ*_, *σ*_*γ*_ ). **(B)** M and V neuron activities over time for one example parameterisation. **(C)** The normalised absolute difference between the averaged mean and the activity of the M neuron (dark green) or between the averaged variance and the activity of the V neuron (brown) for uncorrelated deviations, that is, increasing *σ*_*γ*_ (left), correlated deviations, that is, increasing *μ*_*γ*_ (middle), and the network sparsity. To speed up simulations, we chose *λ*^lower^ = 5 · 10^−2^.

**Figure S3.**
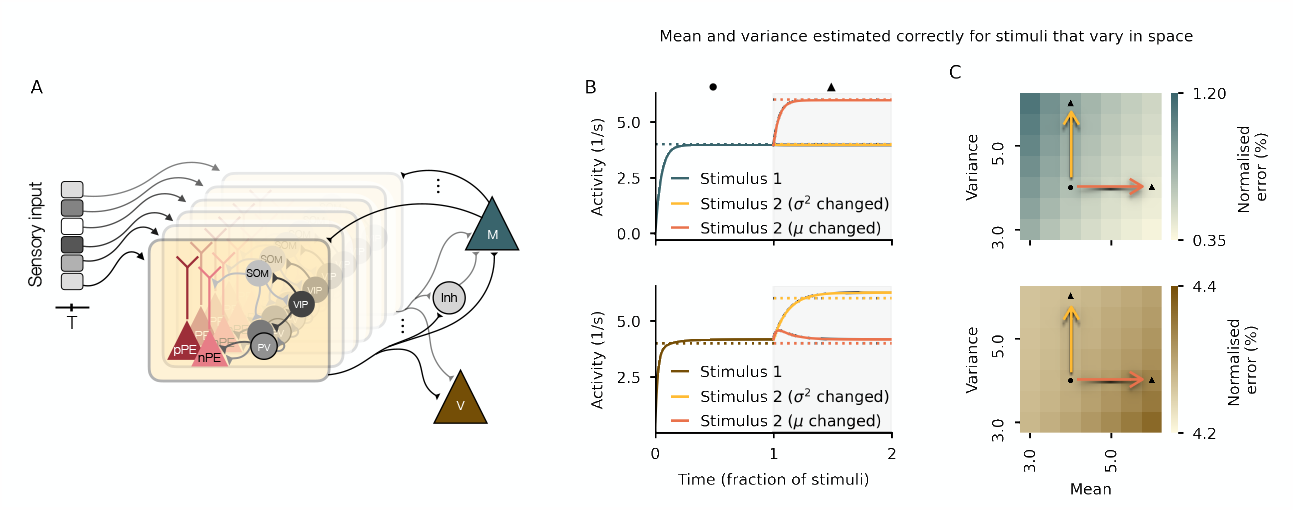
Estimating mean and variance of spatial stimuli. **(A)** Illustration of a network estimating the mean and variance of a stimulus that varies across space. To simulate selectivity, the network comprises 1000 identical, uncoupled mean-field networks each receiving a different input value drawn from a uniform distribution. **(B)** Activity of M neuron (top) and V neuron (bottom) for 2 stimuli. The second stimulus does either differ in the mean (orange) or the variance (yellow) from the first stimulus (indicated in C). **(C)** The normalised absolute difference between the averaged mean and the activity of the M neuron (dark green, top) or between the averaged variance and the activity of the V neuron (brown, bottom) for a range of different stimulus statistics. The examples from B are shown with colored arrows and markers. To speed up simulations, we chose *λ*^lower^ = 3 · 10^−1^.

**Figure S4.**
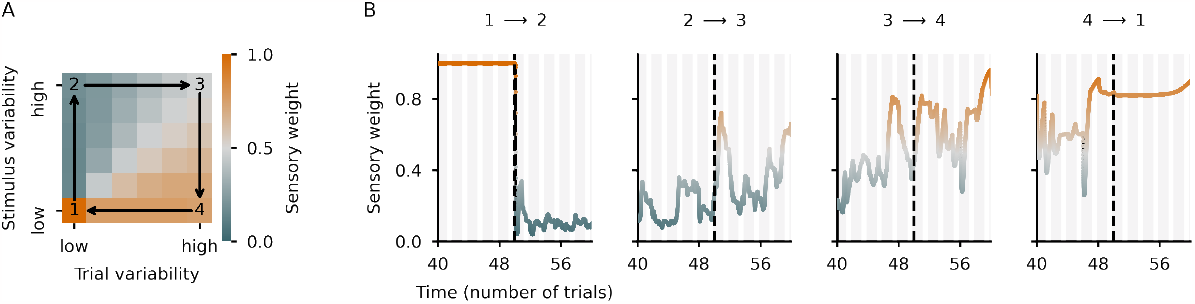
Dynamic variance estimation allows flexible adaptation to changes in the stimulus statistics and environment. **(A)** Illustration of the sensory weight for different input statistics. Numbers denote specific examples. Arrows denote the transitions between those statistics. **(B)** The sensory weight over time is shown for all transitions in (A). For the sake of clarity, we only show the trials 40-60. The switch to new input statistics occurs at trial 50. Parameters are listed in the Supporting Information.

**Figure S5.**
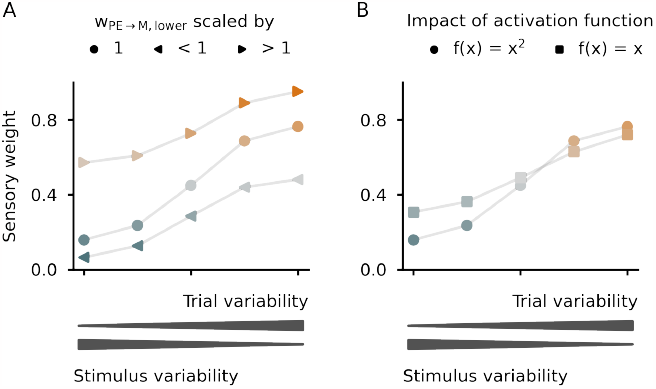
Perturbing the weighting of sensory inputs and predictions by altering network properties. **(A)** The weights from the PE neurons to the M neuron in the lower-order subnetwork are scaled by a factor 0.3 or 7, leading to a distorted sensory weight. If the update of the M neuron in the lower subnetwork is too slow (◂), the prediction is overrated. If the update of the M neuron in the lower subnetwork is too fast (▸), the sensory input is overrated. **(B)** The precise activation function for the V neurons does not have a major impact on the sensory weight. Only for inputs with high stimulus variability, the sensory stimulus is slightly overrated when the quadratic activation function is replaced by a linear, rectified activation function.

**Figure S6.**
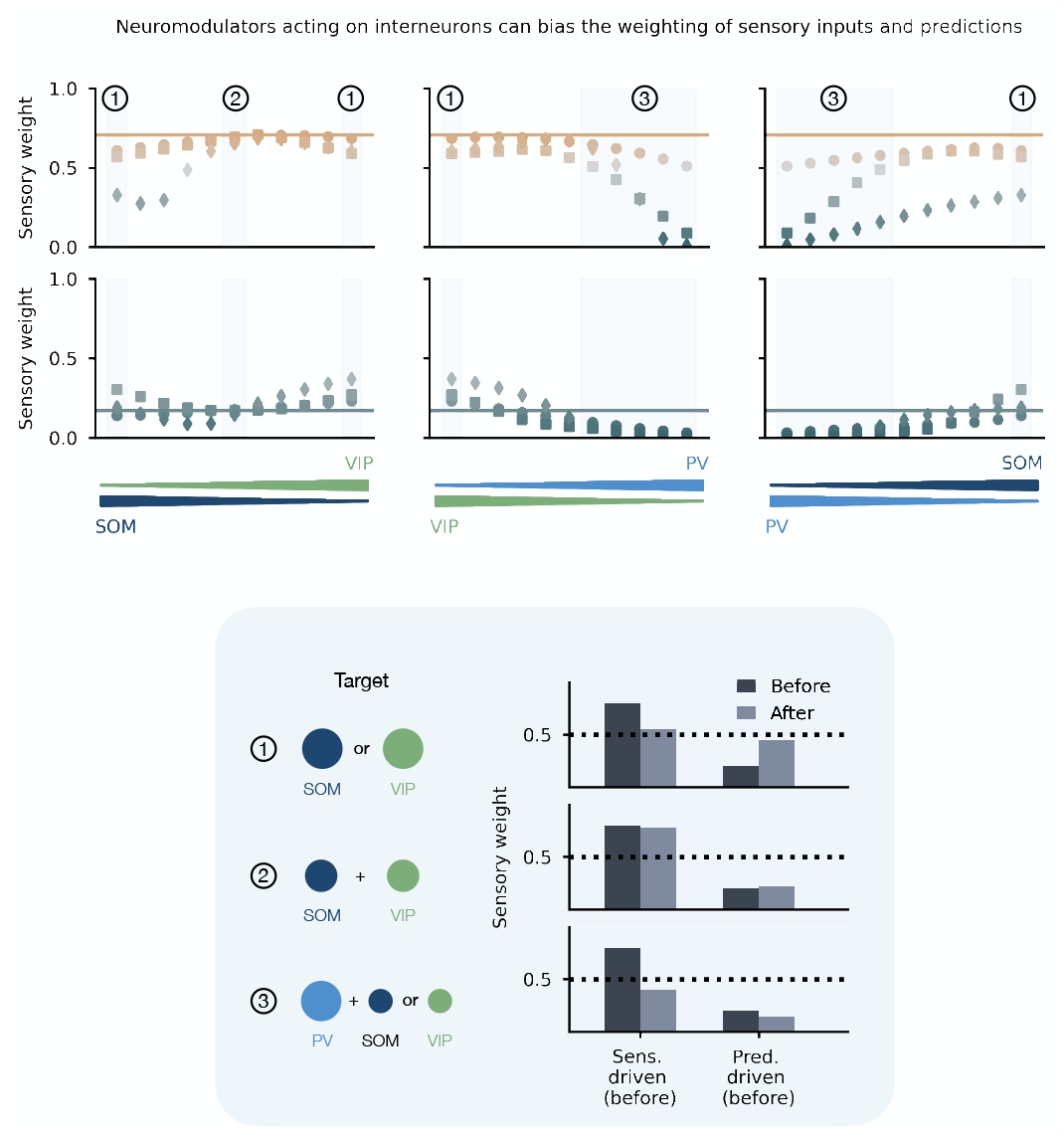
The impact of neuromodulators acting globally on groups of interneurons. The sensory weight changes when groups of interneurons are targeted by a neuromodulator. Whether the sensory weight decreases or increases not also depends on the modulation strength (see Fig. 4) but also on how strongly which interneuron is targeted. As shown in Fig. 4, the sensory weight is pushed toward 0.5 if the VIP neuron is stimulated. The sensory weight generally decreases when PV neurons are the main target. Considered are two limit cases (upper row: more sensory-driven before modulation, lower row: more prediction-driven before modulation). The results are shown for three mean-field networks (see Fig. 4).

**Figure S7.**
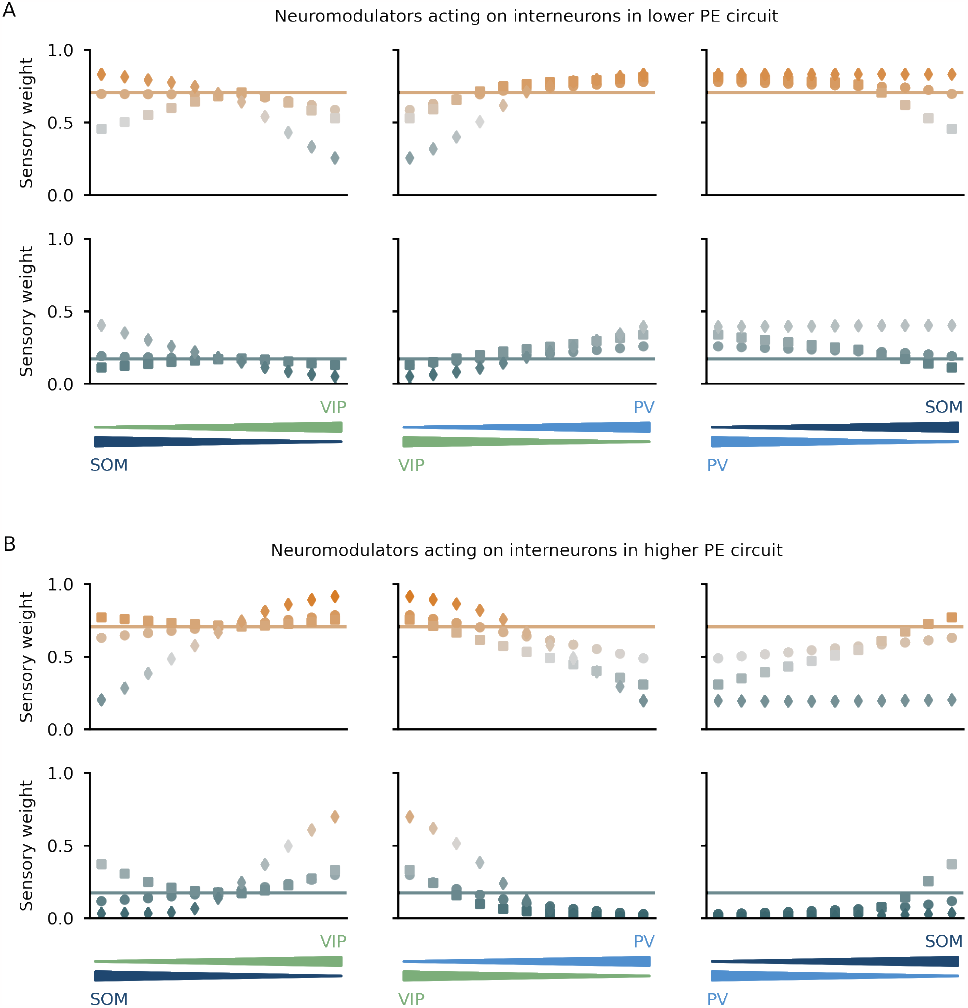
The impact of neuromodulators acting locally on groups of interneurons. **(A)** Sensory weight changes with neuromodulators acting on interneurons in the lower PE circuit. **(B)** Sensory weight changes with neuromodulators acting on interneurons in the higher PE circuit. In general, the changes in the sensory weight is the opposite of the changes seen for neuromodulators acting on the lower-level PE neurons. Simulation parameters, labels and colors as in Fig. 4.

**Figure S8.**
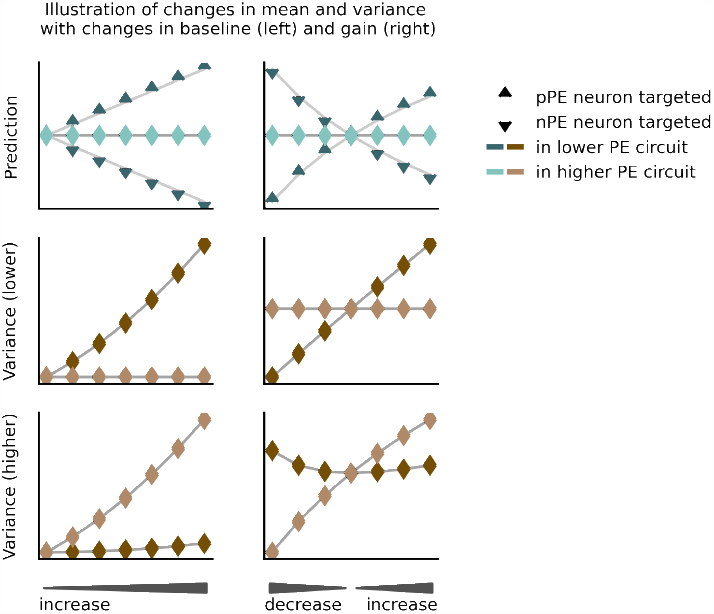
Biased mean and variance estimation by changing the baseline and the gain of nPE and pPE. In a toy model, described in sections B.2 and B.3, the contribution of gain and baseline to the changes in the mean and variance estimation are summarized. The results shown are based on the Eqs. (22), (25), (27) and (30).

**Figure S9.**
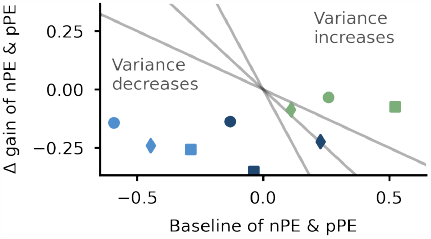
The combined changes in baseline and gain of all PE neurons determine the shift in the sensory weight. Whether and how a neuromodulator changes the sensory weight depends on the interneuron targeted and the effect this interneuron has on the baseline and gain of both PE neurons, which in turn does depend on the network it is embedded in. For small inputs, changes in the baseline dominate, while for large inputs, the changes in the gains dominate the shift in the sensory weight. Gray lines denote different mean inputs, illustrating that the same interneuron can decrease or increase the variance depending on the input regime.

**Figure S10.**
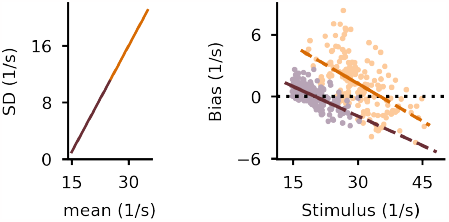
Including scalar variability in the model. When scalar variability is included, that is, the stimulus standard deviation depends linearly on the stimulus mean, the bias is larger for stimuli drawn from the upper end of the stimulus distribution than from the lower end.

